# Extreme mitochondrial reduction in a novel group of free-living metamonads

**DOI:** 10.1101/2023.05.03.539051

**Authors:** Shelby K. Williams, Jon Jerlström Hultqvist, Yana Eglit, Dayana E. Salas-Leiva, Bruce Curtis, Russell J. S. Orr, Courtney W. Stairs, Tuğba N. Atalay, Naomi MacMillan, Alastair G. B. Simpson, Andrew J. Roger

## Abstract

Metamonads are a diverse group of heterotrophic microbial eukaryotes adapted to living in hypoxic environments. All metamonads but one harbour metabolically altered ‘mitochondrion-related organelles’ (MROs) with reduced functions relative to aerobic mitochondria, however the degree of reduction varies markedly over the metamonad tree. To further investigate metamonad MRO diversity, we generated high quality draft genomes, transcriptomes, and predicted proteomes for five recently discovered free-living metamonads. Phylogenomic analyses placed these organisms in a group we informally named the ‘BaSk’ (Barthelonids+Skoliomonads) clade, which emerges as a deeply branching sister group to the Fornicata, a metamonad phylum that includes parasitic and free-living flagellates. Extensive bioinformatic analyses of the manually curated gene models showed that these organisms are predicted to have extremely reduced MRO proteomes in comparison to other free-living metamonads. Loss of the mitochondrial iron-sulfur cluster (ISC) assembly system in some organisms in this group appears to be linked to the acquisition in their common ancestral lineage of a SUF-like minimal system (SMS) Fe/S cluster pathway through lateral gene transfer (LGT). One of the isolates, *Skoliomonas litria*, appears to have undergone further mitochondrial reduction having lost all other known MRO pathways. No proteins were confidently assigned to the predicted MRO proteome of this organism suggesting that the organelle has been lost. The extreme mitochondrial reduction observed within this free-living anaerobic protistan clade is unprecedented and demonstrates that mitochondrial functions, under some conditions, may be completely lost even in free-living organisms.

## Introduction

Many unicellular eukaryote (protist) lineages have independently adapted to low-oxygen conditions. One of their most conspicuous adaptations are highly modified mitochondria referred to as “mitochondrion-related organelles” (MROs) that can function without oxygen. MROs have been best studied in the Metamonada, a supergroup of anaerobic protists with varied lifestyles including parasites, commensals, and free-living marine or freshwater flagellates and amoebae. All metamonad MROs studied to date are functionally reduced compared to aerobic mitochondria with their properties varying markedly across the group (Leger et al., 2017).

The best studied metamonad MROs are hydrogenosomes that occur in the urogenital tract parasite *Trichomonas vaginalis*. The ‘hydrogenosomal’ ATP-producing pathway uses enzymes typically not found in aerobic mitochondria such as pyruvate:ferredoxin oxidoreductase (PFO) to oxidatively decarboxylate pyruvate to acetyl-CoA and reduce ferredoxin, an acetate:succinate CoA transferase that catalyzes the transfer of CoA onto succinate to produce succinyl-CoA, and an iron-only [FeFe]-hydrogenase that reduces protons to hydrogen gas by re-oxidizing the ferredoxin. ATP is produced from ADP and P_i_ by the Krebs cycle enzyme succinyl-CoA synthetase by substrate-level phosphorylation, converting succinyl-CoA to succinate in the process. Trichomonad hydrogenosomes also house a variety of other pathways including a mitochondrial-type iron-sulfur cluster (ISC) biogenesis system, amino acid metabolism, and oxygen detoxification (Schneider et al., 2011; Stairs et al., 2015). Even more highly-reduced MROs called ‘mitosomes’ are found in the metamonad gut parasite *Giardia intestinalis.* These organelles appear to lack any capacity to generate ATP; instead they participate in Fe/S cluster biogenesis via the mitochondrial-type ISC system (Jedelský et al., 2011). Like trichomonads, *Giardia intestinalis* catalyzes acetyl-CoA production from pyruvate using PFO. However, this occurs in the cytoplasm. Acetyl-CoA is converted to acetate and ATP is produced by substrate-level phosphorylation catalyzed by the enzyme ADP-forming acetyl-CoA synthetase (ACS; EC 6.2.1.13) (Sánchez et al., 2000). The most extreme form of MRO reduction in Metamonada can be found in the oxymonad gut commensal *Monocercomonoides exilis* that has lost all traces of the organelle, making it the first truly amitochondriate eukaryote discovered (Karnkowska et al., 2016).

More recent investigations of newly discovered free-living metamonads have revealed novel configurations of MRO functions that do not fit into traditionally-defined organelle classes (Müller et al., 2012). For example, the free-living metamonad *Dysnectes brevis* encodes MRO targeted [FeFe]-hydrogenases and associated proteins but lacks organelle-targeted enzymes for substrate-level phosphorylation, suggesting that while the MRO is capable of hydrogen metabolism, ATP is most likely produced in the cytosol (Leger et al., 2017). A similar hydrogen-producing organelle and predicted cytosolic ATP production was reported in a transcriptomic investigation of the free-living metamonad *Barthelona* sp. PAP020 (Yazaki et al., 2020). Recently, the enigmatic *Anaeramoeba* species – recently-discovered distant relatives to parabasalids like *Trichomonas* – were predicted to possess hydrogenosomes that have retained more mitochondrial systems than any other metamonad studied to date (Stairs et al., 2021). The MRO of the freshwater flagellate *Paratrimastix pyriformis* was recently investigated using spatial proteomic techniques which revealed that the organelle does not participate in either ATP production through extended glycolysis or Fe/S cluster production, and instead produces essential intermediates for the methionine cycle (Zítek et al., 2022). These studies of newly discovered free-living metamonads have provided important insights into the steps by which dramatically distinct MRO-types evolved within the Metamonada.

The multiple independent transitions from aerobically respiring mitochondria to MROs in the eukaryote tree in most cases have involved the complete loss of the mitochondrial genome itself, the loss of aerobic metabolism genes encoded in the nucleus, and the gain of genes encoding novel biochemical pathways by lateral gene transfer (LGT) from bacteria, archaea or other anaerobic protists (Roger et al. 2017). For example, the ‘hydrogenosomal’ metabolism enzymes (Leger et al., 2016; Stairs et al., 2011, 2021), the ability to synthesize rhodoquinone (Stairs et al., 2018) and oxygen defense enzymes (Jiménez-González et al., 2019) all appear to have been acquired by LGT in multiple distinct lineages from bacteria or other protists (Gawryluk & Stairs, 2021; Roger et al., 2017). The most striking examples of the remodeling of mitochondrial functions in anaerobic protists involve the wholesale replacement of the highly-conserved essential mitochondrial ISC system by laterally acquired prokaryotic Fe/S cluster biogenesis systems. In the Archamoebae, a two-protein Fe/S biosynthesis pathway (recently classified as the minimal iron sulfur system (MIS) related to the nitrogen fixation system (Garcia et al., 2022)) was acquired from bacteria and, in free-living members of the group, was subsequently duplicated into cytosolic and MRO-functioning versions (Nývltová et al., 2015; Záhonová et al., 2022). A completely independent replacement of the mitochondrial ISC system occurred in the ancestors of the breviate flagellate *Pygsuia biforma* that acquired an archaeal-type SUF system (Stairs et al., 2014), recently classified as the SUF-like minimal system (SMS) pathway (Garcia et al., 2022). This breviate SmsCB gene (referred to as SufCB in Stairs et al. 2014) in *Pygsuia biforma* was then duplicated to create cytosolic and MRO-targeted SmsCB proteins. In the oxymonads, yet another replacement of the ISC system occurred independently by the lateral acquisition of genes encoding a more complex multi-protein SUF pathway (Karnkowska et al., 2016; Vacek et al., 2018). This enabled the ancestral oxymonad to produce Fe/S clusters in the cytosol, easing the evolutionary pressure to keep the canonical ISC system, paving the way for the complete loss of MROs in *Monocercomonoides* (Karnkowska et al., 2016). Phylogenetic analyses revealed that this SUF system is shared by a variety of preaxostylans and is likely bacterial in origin (Vacek et al., 2018).

To further investigate the patterns and process of genome and MRO evolution in metamonads, we sequenced the genomes of five recently-isolated free-living bacterivorous flagellates. Three of these organisms, the ‘skoliomonads’ *Skoliomonas litria, Skoliomonas* sp. GEM-RC (GEMRC) and *Skoliomonas* sp. TZLM3-RCL (RCL) were isolated from sediments of alkaline hypersaline soda lakes (for detailed descriptions see Eglit et al., 2024) (Figure 1A-C). The other two are shallow marine-sediment dwelling ‘barthelonids’ (Yazaki et al. 2020): *Barthelona* sp. strain PCE (Figure 1D) and *Barthelona* sp. PAP020 (Yazaki et al., 2020). Unlike most previous studies of this kind that rely on incomplete transcriptomic data, we generated high quality contiguous genome assemblies, transcriptomes, and predicted proteomes from four of these five organisms. Our phylogenomic analyses demonstrate that these five metamonads form a highly supported clade, referred to as ‘BaSk’ (short for ‘Barthelonids+Skoliomonads’), that emerges as a sister group to Fornicata *sensu stricto* (diplomonads, retortamonads and *Carpediemonas*-like organisms (CLOs)). Genome size and coding capacity varies considerably across these organisms, with one possessing the smallest genome of a free-living eukaryote currently known. Their anaerobic metabolisms and MRO functional capacities also vary considerably. Unexpectedly, we found yet another instance of the horizontal acquisition of the SmsCB gene in their common ancestor. While several barthelonids possess both the SmsCB gene and core parts of a mitochondrial-type ISC system, all traces of the ISC system have been lost in the skoliomonads. For *Skoliomonas litria*, no MRO proteins could be confidently identified, suggesting it may lack the organelle altogether.

**Figure 1.**
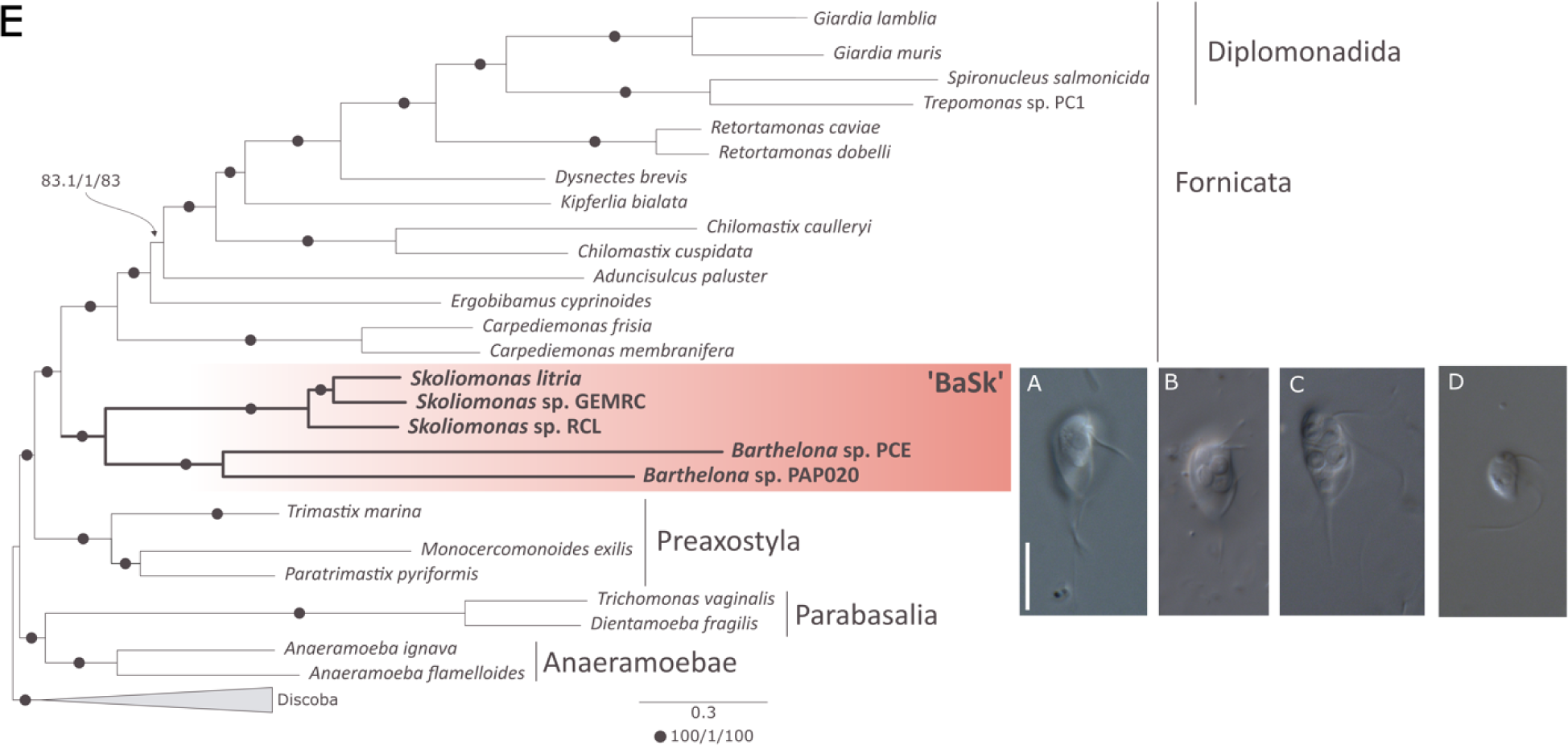
‘BaSk’ is a clade of anaerobic protists that branches as sister to all known Fornicata within the Metamonada. Panels A-D) Differential interference contrast light micrographs of skoliomonad lineages Skoliomonas litria (A), Skoliomonas sp. GEMRC (B), and Skoliomonas sp. RCL (C), as well as Barthelona sp. PCE (D) showing overall morphology (imaging and methods described in Eglit et al. 2024). Scale bars indicate a length of 10 µm. (E) A maximum likelihood (ML) phylogeny based on 174 concatenated aligned proteins encompassing 46,113 sites. The depicted topology was estimated using IQ-TREE under the LG+C60+F+Γ model and was used as a guide tree for estimating the LG+PMSF(C60)+F+Γ model. Support under the latter model was evaluated by SH-aLRT support percentage, aBayes support and nonparametric bootstrap percentage based on 100 nonparametric bootstrap replicates. The scale bar indicates the expected number of amino acid substitutions per site under the PMSF model. When at least one support value was less than maximal, all three are shown on the branch in the order SH-aLRT support percentage/aBayes/nonparametric bootstrap percentage. A black dot on the branch indicates all three approaches yielded full (i.e. 100/1/100) support. The BaSk clade is highlighted in red.

## Results & Discussion

### Near-complete draft genomes of four novel metamonads

Near-complete genome assemblies of *Barthelona* sp. PCE, *Skoliomonas litria*, *Skoliomonas* sp. RCL and *Skoliomonas* sp. GEMRC were generated using a combination of Oxford Nanopore long-read sequencing for assembly and Illumina short-read sequencing for error-correction post-assembly. These assemblies were thoroughly decontaminated using the Eukfinder workflow (Zhao et al., 2023).

To assess the quality of these assemblies we used Merqury, a k-mer counting tool that using genomic Illumina reads, assesses the completeness of the four *de novo* long-read assemblies (Rhie et al., 2020). Based on the Merqury analyses, all four assemblies exceeded 98% completion. Genome completeness was further supported by the fact that, for each of the four organisms, >99% of their respective eukaryotic-classified RNAseq reads mapped to the corresponding genome assembly. The combination of long and short-read data afforded assemblies that were high coverage and comprised of a few relatively large contigs (see Table 1). A draft genome assembly of *Barthelona* sp. PAP020 was also generated using only Illumina reads (Table 1). This relatively fragmented assembly was included with other metamonads considered in this study to supplement our analyses concerning MRO reconstruction, metabolic pathway inventory, and phylogenomic inference, discussed below.

**Table 1.**
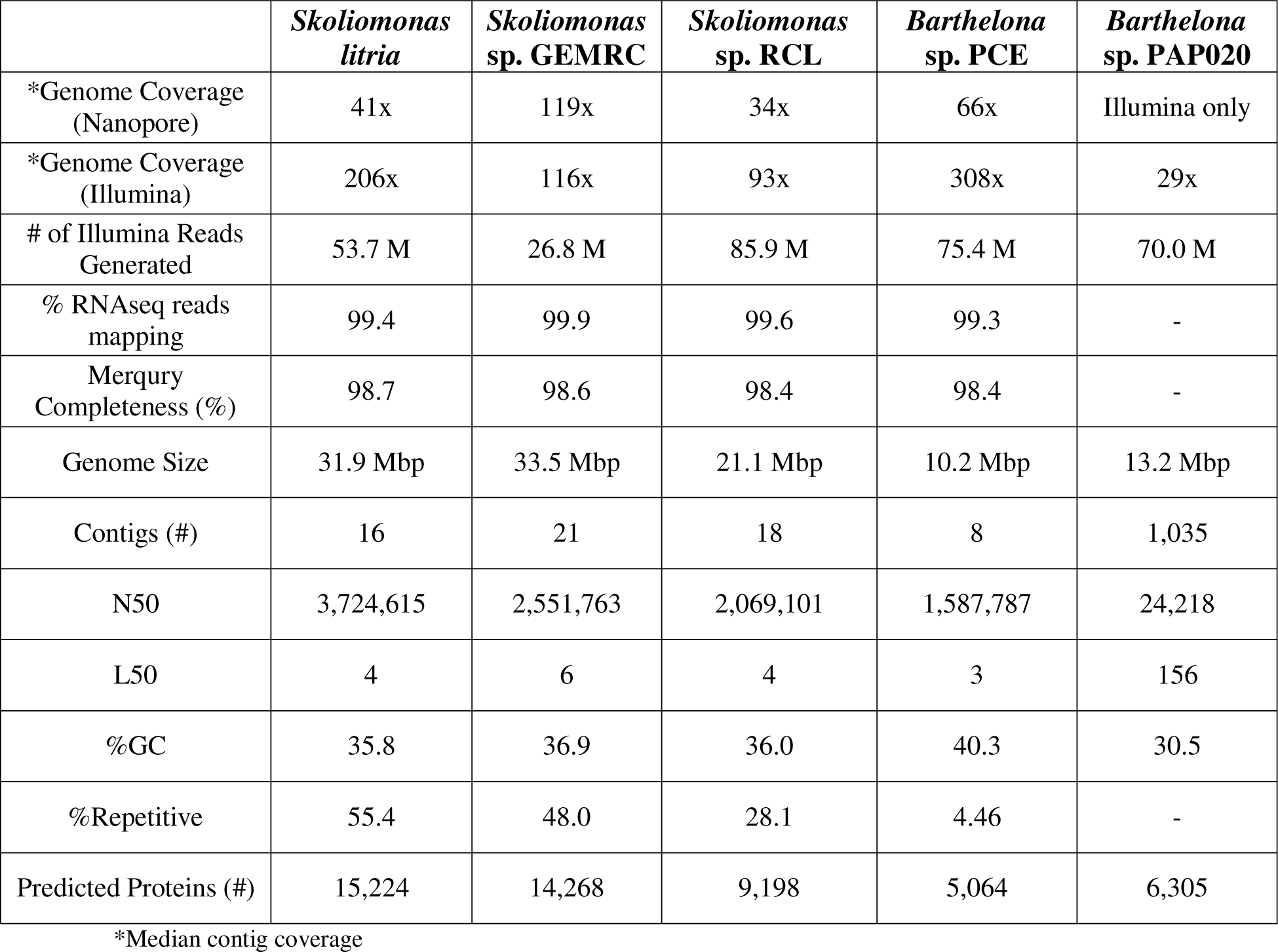
Summary statistics for the genomes, transcriptomes, and predicted proteomes of the BaSk clade.

Gene predictions for these assemblies were generated by an in-house pipeline. To compare the qualities of the various gene predictions, BUSCO analyses were conducted based on the whole genomes themselves, the gene predictions and the transcriptome assemblies (Table 2). To assess potential improvement of the predicted protein set after curation, the gene predictions for *Skoliomonas litria* were further manually curated (Table 2). The percent of BUSCO genes identified amongst the gene predictions (i.e. predicted protein set) ranged from 63.7% (complete+fragmented) for *Barthelona* sp. PCE to 77.9% (complete+fragmented) for *Skoliomonas* sp. RCL. In general, the BUSCO scores of the gene predictions are very similar to those from the transcriptomes but the BUSCO genes of the former are less fragmented. The manual curation of *Skoliomonas litria* predicted proteins increased the reported BUSCO number by only one, indicating that the manual curation did not substantially improve the gene models. Note that these relatively low BUSCO scores are comparable to other well-characterized metamonads assemblies which tend to have highly divergent or missing orthologs relative to most other eukaryotes (Salas-Leiva et al., 2021; Stairs et al., 2021).

**Table 2.**
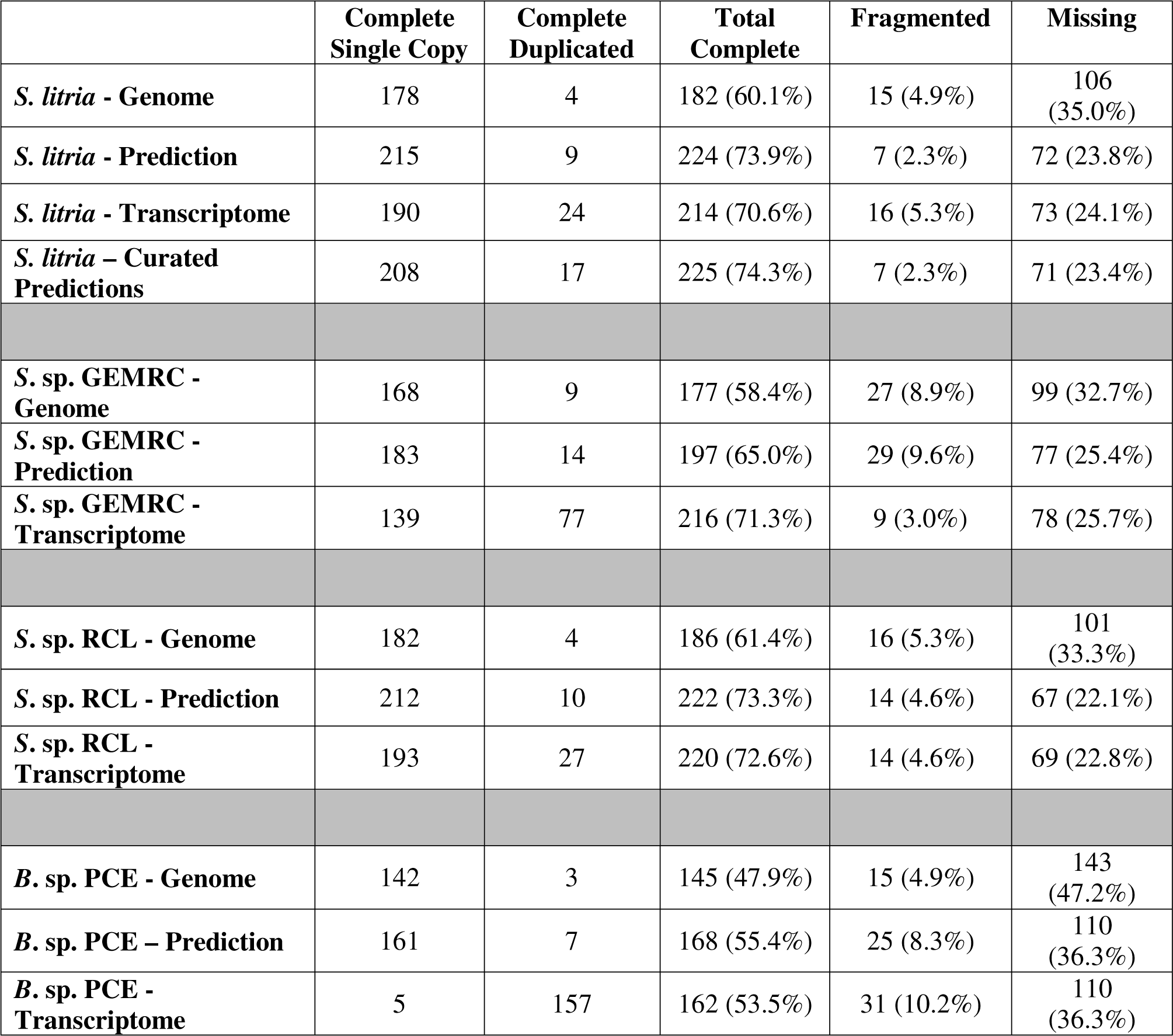
BUSCO scores for members of the BaSk clade.

|  | **Complete**<br>**Single Copy** | **Complete**<br>**Duplicated** | **Total**<br>**Complete** | **Fragmented** | **Missing** |
| --- | --- | --- | --- | --- | --- |
| *S. litria* - Genome | 178 | 4 | 182 (60.1%) | 15 (4.9%) | 106<br>(35.0%) |
| *S. litria* - Prediction | 215 | 9 | 224 (73.9%) | 7 (2.3%) | 72 (23.8%) |
| *S. litria* - Transcriptome | 190 | 24 | 214 (70.6%) | 16 (5.3%) | 73 (24.1%) |
| *S. litria* – Curated<br>Predictions | 208 | 17 | 225 (74.3%) | 7 (2.3%) | 71 (23.4%) |
| *S. sp. GEMRC* -<br>Genome | 168 | 9 | 177 (58.4%) | 27 (8.9%) | 99 (32.7%) |
| *S. sp. GEMRC* -<br>Prediction | 183 | 14 | 197 (65.0%) | 29 (9.6%) | 77 (25.4%) |
| *S. sp. GEMRC* -<br>Transcriptome | 139 | 77 | 216 (71.3%) | 9 (3.0%) | 78 (25.7%) |
| *S. sp. RCL* - Genome | 182 | 4 | 186 (61.4%) | 16 (5.3%) | 101<br>(33.3%) |
| *S. sp. RCL* - Prediction | 212 | 10 | 222 (73.3%) | 14 (4.6%) | 67 (22.1%) |
| *S. sp. RCL* -<br>Transcriptome | 193 | 27 | 220 (72.6%) | 14 (4.6%) | 69 (22.8%) |
| *B. sp. PCE* - Genome | 142 | 3 | 145 (47.9%) | 15 (4.9%) | 143<br>(47.2%) |
| *B. sp. PCE* – Prediction | 161 | 7 | 168 (55.4%) | 25 (8.3%) | 110<br>(36.3%) |
| *B. sp. PCE* -<br>Transcriptome | 5 | 157 | 162 (53.5%) | 31 (10.2%) | 110<br>(36.3%) |

The sizes of the draft genomes and the number of proteins predicted differ markedly between taxa (Table 1). Notably, at 10.2 Mbp, *Barthelona* sp. PCE appears to possess the smallest genome for a free-living eukaryote currently known. It is smaller than the compact genomes of the acidophilic alga *Galdieria sulphuraria* (13.7 Mbp, see Schönknecht et al., 2013), *Saccharomyces cerevisiae* (12.07 Mbp, see Goffeau et al., 1996), and the genome of the smallest known free-living eukaryote, the marine alga *Ostreococcus tauri* (12.56 Mbp, see Derelle et al., 2006). Genome reduction is well known to occur in parasitic and endosymbiotic microbes (Manzano-Marín & Latorre, 2016; Xu et al., 2020), as well as in free-living organisms, where it is thought that compact genomes are an efficiency adaptation to nutrient-poor environments (Giovannoni et al., 2014). However, for heterotrophic phagotrophic protists, like *Barthelona* sp. PCE, nutrients are obtained predominantly from their bacterial prey. It is unclear whether the foregoing generalizations hold in this case or what other forces led to genome reduction in the barthelonid lineage.

Two of the genome assemblies have apparent telomeric repeats on the ends of some contigs. *Barthelona* sp. PCE possesses two contigs with what appear to be telomeres at one end of the contig with an unusual telomere repeat sequence of 5’-TATATGGTCT-3’. Although this telomeric sequence is quite different from the eukaryotic ‘consensus’ repeat, other protists across the eukaryote tree of life have also been reported to have highly divergent telomeric repeats (see Fulnečková et al., 2013); the significance of these unusual telomeres is unknown. The *Skoliomonas* sp. GEMRC assembly has six telomere-to-telomere assembled chromosomes and an additional nine contigs with telomeres at only one end of the contig. The latter telomeres have the canonical 5’-TTAGGG-3’ repeating sequences.

### The BaSk taxa are the sister group to known members of Fornicata

To determine the phylogenetic affinities of the BaSks within the metamonads, we added orthologs from the five draft assemblies to a previously constructed phylogenomic data set of orthologous proteins (Brown et al., 2018; S. Kang et al., 2017). After single-protein alignment manual curation to remove contaminants and paralogs, the final dataset of 174 highly conserved proteins was analyzed by maximum likelihood under the site-heterogeneous model LG+C60+F+ Γ (see Figure 1E). The resulting tree confirms the placement of *Barthelona* sp. PAP020 as previously reported (Yazaki et al., 2020) and placed barthelonid strain *Barthelona* sp. PCE as its immediate sister. The skoliomonads form a clade with the barthelonid group, with *Skoliomonas litria* and *Skoliomonas* sp. GEMRC branching together to the exclusion of *Skoliomonas* sp. RCL. The ‘BaSk clade’ (Barthelonids + Skoliomonads) are sister to fornicates with maximum support (i.e. both the fornicate group and the fornicate + BaSk clade are supported with 100% non-parametric bootstrapping support, aBayes support, and approximate likelihood-ratio test support, all evaluated under the PMSF model).

Members of Fornicata possess a particular cytoskeletal feature, a distinct fibrous arch, that defines the group (Simpson, 2003). As there are no ultrastructural data for the flagellar apparatus cytoskeleton of any BaSk member to date, it is unclear whether they should be classified within the Fornicata or remain as a distinct sister group. Regardless, the BaSks ‘deep’ phylogenetic placement within the metamonads make them a key group for investigations into the origins and evolutionary trajectories of genes involved in parasitism and MRO reduction within the Metamonada supergroup.

### BaSks all possess highly reduced MROs and share the same core anaerobic metabolism

We searched the genomes, transcriptomes, and predicted protein sets of all four BaSks using a variety of bioinformatic approaches (see Materials and Methods) to recover proteins related to MRO function and anaerobic metabolism. We also utilized a draft short-read Illumina genome assembly of *Barthelona* sp. PAP020 in these searches to supplement our comparisons. Several mitochondrial targeting sequence prediction programs were used to estimate the probability of mitochondrial localization of each predicted protein in each of the five genomes. We used these predicted proteins as queries against the nr database using BLAST and looked for outputs relating to mitochondrial localizing proteins. The predicted proteins from the BaSk genomes were also used as queries in a BLAST search of a database comprised of mitochondrial proteomes from six diverse eukaryote lineages and the best hits from each query were retained and inspected. Next, we utilized datasets of known metamonad MRO proteins to query the predicted protein sets from the BaSk genomes, as well as directly investigate the genomes and transcriptomes. This was done through both BLAST search and HMM profile searching. In cases where search results were ambiguous, we utilized appropriate Pfam profiles to search for protein domain information. Finally, we assessed candidate proteins by constructing phylogenetic trees of corresponding MRO proteins with additional cytosolic homologs, to further verify the probable localization of BaSk proteins. A summary of the results of all of these searches can be found as Figure 2. Below we discuss the systems we investigated in detail by first highlighting the pathways that, when present, function exclusively in MROs and are therefore strong indicators of the presence of the organelle.

**Figure 2.**
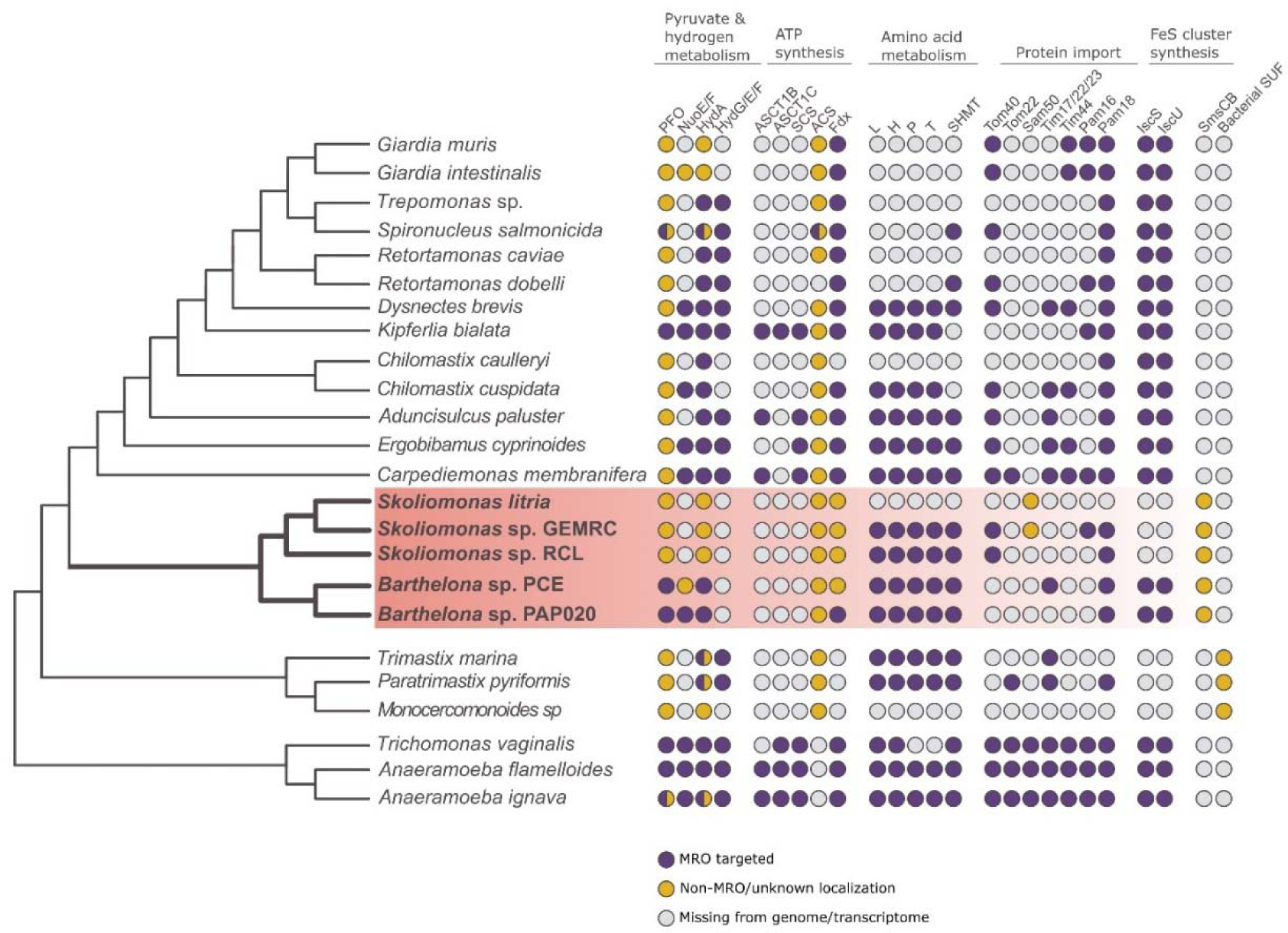
Presence, absence and predicted localization of key MRO and energy metabolism enzymes in ‘BaSks’ and other metamonads. A schematic phylogeny of the Metamonada is shown on the left based on Fig. 1, with the BaSk clade highlighted in red. The pathways depicted include anaerobic extended glycolysis/energy metabolism enzymes, conserved MRO proteins, MRO protein targeting machinery components, and the SmsCB fusion protein. Coloured versus grey circles indicate presence/absence of the protein from predicted proteomes and/or transcriptome data and the colours indicate probable localization. Circles split in half indicate that more than one paralog was present with different predicted localizations. Protein abbreviations: PFO - Pyruvate:ferredoxin oxidoreductase; NuoE/F - Respiratory-chain NADH dehydrogenase 24/51 kDa subunit; HydA – Iron-only hydrogenase; ASCT1B – Acetate:succinyl-CoA transferase B; ASCT1C - Acetate:succinyl-CoA transferase C; SCS - Succinyl-CoA synthase; ACS - Acetyl-CoA synthase; Fdx – Ferredoxin; L – Glycine cleavage system subunit L; H - Glycine cleavage system subunit H; P - Glycine cleavage system subunit P; T - Glycine cleavage system subunit T; SHMT - Serine hydroxymethyltransferaseTom40 - Translocase of outer mitochondrial membrane 40; Tom22 - Translocase of outer mitochondrial membrane 22; Sam50 – Sorting and assembly machinery 50; Tim17/22/23 - Translocase of the inner membrane 17/22/23; Tim44 - Translocase of the inner membrane 44; Pam16 - Presequence translocase-associated motor 16; Pam18 - Presequence translocase-associated motor 18; IscS - Iron-sulfur cluster assembly enzyme cysteine desulfurase; IscU - Iron-sulfur cluster assembly enzyme scaffold; SufCB - Sulfur formation CB fusion protein.

We first investigated whether the BaSk genomes encoded homologs of the glycine cleavage system (GCS), an exclusively mitochondrial pathway which converts glycine to ammonia and carbon dioxide while also reducing NAD+ to NADH and generating 5,10-methylene-tetrahydrofolate (5,10-CH_2_-THF) (Kikuchi, 1973). This pathway is commonly retained in MROs of free-living anaerobic protists including some metamonads (Leger et al., 2017), but is sometimes missing in parasitic lineages (reviewed in Roger et al., 2017). Except for *Skoliomonas litria*, all BaSks encode the complete GCS (i.e. the T-, P-, L- and H-proteins). Additionally, all BaSks possessing the GCS also have a single serine hydroxymethyltransferase (SHMT) gene. This enzyme produces serine using both glycine and the 5,10-CH_2_-THF produced by the GCS. GCS and SHMT are found in the MROs of a variety of diverse free-living metamonads including *Paratrimastix pyriformis* (Zítek et al., 2022; Zubáčová et al., 2013) and all of the non-diplomonad fornicates (i.e. CLOs and *Chilomastix cuspidata*; see Leger et al., 2017). Complete absence of the GCS in metamonads is seen in the parasitic diplomonads *Giardia intestinalis* and *Spironucleus salmonicida* (Jerlström-Hultqvist et al., 2013; Leger et al., 2017). Other parasitic metamonads such as *Trichomonas vaginalis* retain some of the GCS subunits (Mukherjee et al., 2006), but have repurposed them to function in a peroxide detoxification pathway (Nývltová et al., 2016). It was particularly unexpected, therefore, that we found neither genes encoding GCS or SHMT in *Skoliomonas litria,* making it the only known free-living metamonad completely lacking this pathway, and the only BaSk member to have no detectable MRO amino acid metabolism. The MROs within the BaSks are therefore key to understanding the reduction of amino acid metabolism that is usually only ever seen in parasitic metamonads. As previously reported (Yazaki et al., 2020), *Barthelona* sp. PAP020 contains a OsmC homolog, but we were unable to detect this enzyme in any of the other BaSk data, including the *Barthelona* sp. PAP020 genome. Because this enzyme lacks a mitochondrial targeting sequence (MTS), it is unclear where this protein localizes in *Barthelona* sp. PAP020, and if it participates in peroxide detoxification in the *Barthelona* sp. PAP020 MRO, as it does in the MRO of *Trichomonas* (Nývltová et al., 2016).

Next, we investigated the highly conserved mitochondrial Fe/S cluster (ISC) pathway that is also exclusively found within mitochondria or MROs. Unexpectedly, amongst all the BaSks, only the barthelonid PCE and PAP020 assemblies encoded *any* detectable ISC system components. These two genomes encode cysteine desulfurase NFS1 (also known as ISCS), the scaffold protein ISCU and a ferredoxin. Notably, they also encode Fe/S cluster containing NuoE and NuoF subunits of complex I, which, like core ISC components, are homologs which are only known to function in mitochondria and the MROs of other organisms. Surprisingly, with the exceptions of mitochondrial Hsp70 (discussed further below) and ferredoxin, no other genes encoding ISC pathway components were detected in the barthelonids or any other of the BaSks (see Supplementary Table 1 for ISC proteins queried). The absence of some of these proteins, like Isd11 and ferredoxin reductase (Arh1), is not unexpected as they have not been detected in most metamonads (Motyčková et al., 2023). The absence of other early ISC pathway proteins implicated in [2Fe-2S] cluster synthesis (e.g. frataxin, Jac1 (HscB) and Mge1 (GrpE)) is more puzzling as these components are usually detected in metamonads (Motyčková et al., 2023). Similarly, some late ISC pathway components implicated in [4Fe-4S] cluster synthesis (Lill & Freibert, 2020) are found in other metamonads (e.g Grx5, Isa2, NFU1 and BolA; Motyčková et al., 2023), but were not detected in any BaSks (Supplementary Table 1). This is strange given that the barthelonids possess mitochondrial NuoF subunits that typically house an [4Fe-4S] cluster (Ohnishi, 1998). This means that if these NuoF subunits are indeed found within the MROs of the barthelonids, as they are with other eukaryotes, their [4Fe-4S] clusters are being produced by an unknown mechanism.

Despite the lack of ISC system proteins in most BaSks, all BaSk members appear to encode a similar partial cytosolic CIA system that includes apoprotein targeting components CIA1, CIA2, and NAR1, the NBP35 scaffold protein, and TAH18, which aids in maturation of Fe/S cluster proteins. This reduced CIA pathway is not unlike the CIA pathway found within *Giardia intestinalis* (Pyrih et al., 2016).

### BaSks possess simple archaeal-type SUF-like minimal system (SMS) fusion proteins

We investigated the possibility that the lack of ISC system components in BaSk members may be complemented by the presence of an alternative Fe/S cluster system such as the minimal Fe/S (MIS) or sulfur mobilization (SUF) systems found in a number of other anaerobic protists (Anwar et al., 2014; Garcia et al., 2022; Stairs et al., 2014; Vacek et al., 2018; Žárský et al., 2021). In all of the BaSk genomes, we were able to identify a simple archaeal-type SUF-like minimal system (SMS) consisting of a gene encoding an SmsCB fusion protein that was most similar to the simple archaeal-type SmsCB systems found in the breviate *Pygsuia biforma* (Stairs et al., 2014), the anaerobic jakobid *Stygiella incarcerata* (Leger et al., 2016) and gut commensal opilinatan stramenopile *Blastocystis* (Tsaousis et al., 2012; Yubuki et al., 2020). To investigate the origin of this protein, we performed phylogenetic analysis separately for the SmsC and SmsB domains (Supplementary Figures 1 & 2) before concatenating the domains together for the final tree (Figure 3). The BaSk SmsCB proteins form a strongly supported clade that is sister group to all archaeal-type SmsCB fusion proteins from anaerobic protists. Consistent with previous reports, the anaerobic protistan clade of SmsCB fusion proteins is sister group to the orthologs from members of the Methanomicrobiales order of Archaea (Tsaousis et al., 2012). This phylogenetic pattern is most easily explained if there were an original transfer and fusion of SmsCB to an anaerobic protist from a methanomicrobiales donor, followed by a series of eukaryote-to-eukaryote LGT events amongst disparate lineages of anaerobic eukaryote lineages. The deeply branching position of the BaSk clade amongst the anaerobic protists suggests it is possible that an ancestral BaSk was the first recipient of these genes and an early offshoot from the BaSk lineage passed the fusion gene to other anaerobic protists, although it is impossible to rule out alternative scenarios. Regardless, this SMS system is phylogenetically distinct, and has a separate origin from, the more complex SUF system found within the metamonad preaxostylans (i.e. oxymonads and *Paratrimastix pyriformis*) (Vacek et al., 2018) and therefore represents an interesting case of convergence by LGT within the Metamonada. The resulting reduction and loss of the ISC system following acquisition of a SMS system is also similar between these two lineages and the breviate *Pygsuia biforma*. The gain of an alternate Fe/S cluster synthesis system in these cases likely ‘preadapts’ the ancestral organism to the loss of the ISC system in some of the lineages (e.g. an ancestor of *Skoliomonas* sp. GEMRC, *Skoliomonas* sp. RCL and *Skoliomonas litria*). Unlike *Pygsuia biforma* and the preaxostylans, *Barthelona* sp. PCE and *Barthelona* sp. PAP020 retain a few core components of the early ISC system in addition to possessing the novel SMS system, suggesting that their MRO ISC system is at least partially functional. This retention of the ISC proteins may be related to the possession of Fe/S cluster-containing proteins such as the NuoE and NuoF complex I proteins that are predicted to function in the MRO. Although we cannot definitively determine the subcellular localization of the SmsCB in BaSks bioinformatically, it is likely that this system is cytosolic as in other anaerobic protists such as *Blastocystis* spp. (Tsaousis et al., 2012). If it is indeed cytosolic, it is unclear whether it would function in coordination with the cytosolic CIA Fe/S biogenesis system. One possibility is that the CIA system and the SmsCB system may function in parallel with each responsible for the assembly of Fe/S clusters for different specific subpopulations of apoproteins. Alternatively, the two systems may work together, similar to how the ISC system coordinates with the CIA pathway (Pyrih et al., 2016). In the latter scenario, the SmsCB fusion protein would provide the CIA pathway with a sulfur-containing intermediate, which the CIA pathway would then use to assemble Fe/S clusters and insert them into recipient apoproteins.

**Figure 3.**
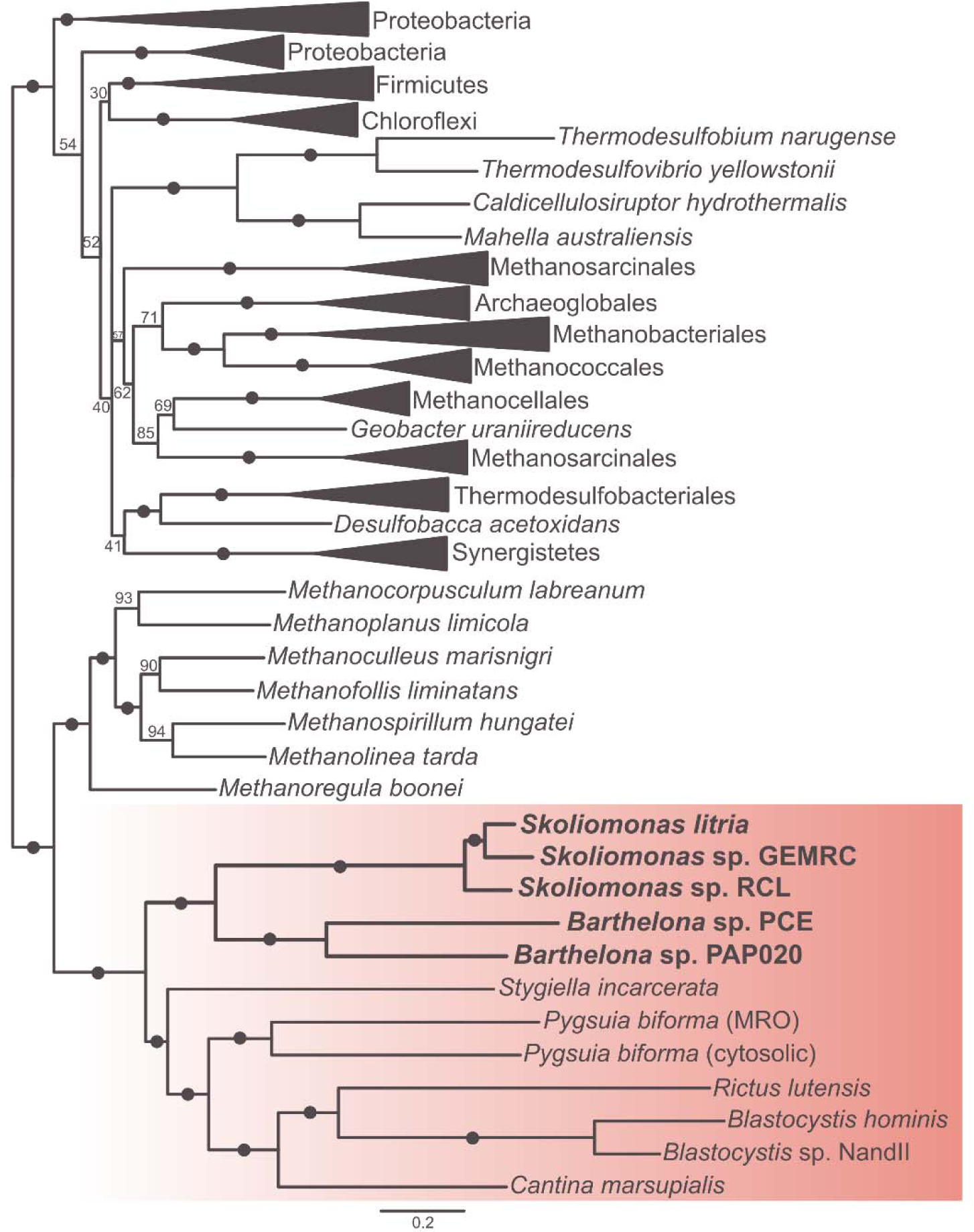
A phylogeny of the SmsCB fusion protein in a variety of anaerobic eukaryotes (red) and their closest bacterial and archaeal homologs. The alignment contains the SmsCB fusion protein from the eukaryotes and concatenated SmsC and SmsB proteins from the prokaryotes. This ML tree was estimated under the LG+C60+F+Γ model using IQ-TREE. Ultrafast bootstrap values are displayed on the branches, with dots indicating UFBOOT value over 95%.

In any case, the skoliomonads completely lack any dedicated ISC components and do not possess proteins with identifiable cysteine desulfurase domains as assessed by HMM profile searches and phylogenetic analysis (see Supplementary Figure 3). This is surprising because for most Fe/S cluster synthesis systems, cysteine is the source of sulfur and a cysteine desulfurase is the key enzyme involved in mobilizing it. We suspect that the SmsCB system of these organisms may directly utilize sulfide from the environment as the sulfur source for Fe/S clusters, as previously suggested for *Pygsuia biforma* (Stairs et al., 2014). Indeed, direct sulfide utilization has been demonstrated for methanogenic archaea that possess a simple cysteine-desulfurase-lacking SMS pathway and which also live in sulfidic conditions (Liu et al., 2010). The exact mechanism by which such organisms construct Fe/S clusters is unknown and more work needs to be done to understand their biogenesis in these organisms.

### Presence or absence of other conserved mitochondrial proteins in the BaSk taxa

The next highly conserved mitochondrion/MRO proteins that we investigated were the mitochondrial orthologs of the molecular chaperones chaperonin-60 (Cpn60) and hsp70. To distinguish cytosolic Chaperonin Containing TCP-1 (CCT) subunits from mitochondrial Cpn60 (GroEL) homologs and cytosolic and endoplasmic reticulum (ER) paralogs of hsp70 from their mitochondrial Hsp70 (i.e. dnaK orthologs), we used phylogenetic analyses. For *Skoliomonas litria* and *Skoliomonas* sp. RCL, all Cpn60 candidates identified distinctly branched with CCT subunit clades (See Supplementary Figure 4); no bona fide mitochondrial Cpn60s could be identified from these species. *Skoliomonas* sp. GEMRC, *Barthelona* sp. PCE, and *Barthelona* sp. PAP020 appear to have both CCTs and mitochondrial Cpn60. Phylogenetic analysis of Hsp70 proved more difficult, due in part to a greater degree of similarity between paralogous copies of the chaperone. To supplement this analysis, signature insertion-deletions in the multiple alignments were identified that were used in conjunction with well-annotated Hsp70 yeast homologs (Nelson et al., 1992) to group sequences into cytosolic/ER versus mitochondrial type Hsp70 groups (see Supplementary Figure 5). This information was then checked against the phylogeny for subcellular localization predictions of BaSk Hsp70 candidates. From this analysis, mitochondrial-type Hsp70s were identified in all BaSks except *Skoliomonas litria*. Within MROs, Cpn60 and Hsp70 aid in protein translocation into the matrix and help newly translocated proteins adopt their native structure. These chaperones are ubiquitously predicted to function in a variety of MROs, so it is notable that they are missing in some of the BaSks data. For *Skoliomonas litria*, we did not detect any genes encoding MRO chaperones, while *Skoliomonas* sp. RCL appears to contain Hsp70 but lacks Cpn60. This further highlights the highly reduced nature of the MROs found within these protists, especially within *Skoliomonas litria*.

All mitochondria and MROs have membrane associated proteins, including the TIM/TOM complex proteins involved in protein translocation into the organelle. To search for these, we used HMM profiles as many of these sequences are highly divergent and not detectable by pairwise alignment-based search algorithms. The main channel forming protein in the TOM complex, Tom40, was only identifiable in *Skoliomonas* sp. RCL and *Skoliomonas* sp. GEMRC, while the receptor Tom70 was identified in *Barthelona* sp. PCE, *Barthelona* sp. PAP020, and *Skoliomonas* sp. RCL. TIM subunits were generally absent except for Tim17 that was identified in *Barthelona* sp. PCE only. Except for *Skoliomonas litria*, all BaSk members appeared to have the co-chaperone Pam18 subunit of the presequence translocase-associated motor (PAM) complex in addition to the core mtHsp70 subunit. Pam16 was only identified in *Skoliomonas* sp. GEMRC and Pam17 appeared to be absent.

In mitochondria, mitochondrial processing peptidases (MPPs) and presequence proteases (PreP) remove the MTS after protein import into the organelles (Garrido et al., 2022). We identified MPP/PreP homologs only in *Skoliomonas litria*, *Skoliomonas* sp. GEMRC, and *Skoliomonas* sp. RCL and none had predicted MTSs. We further investigated their domain structures and phylogenetic position amongst MPP/PreP-related protein families in eukaryotes and prokaryotes. For our phylogenetic analysis we used a recently published multiple alignment of all major families of M16 proteases (Garrido et al., 2022) supplemented with top 20 BLAST hits in the nr database to each of the *Skoliomonas litria* sequences. Our phylogenetic analyses (Supplementary Figure 6A) showed that skoliomonad sequences branch in multiple different positions in the tree, but none are closely related to eukaryote mitochondrial MPPs or PrePs. All but one of these proteins have a four-domain structure that is characteristic of PreP and SPPs but distinct from the two domain structure of the MPP family (Garrido et al., 2022). One *Skoliomonas* sp. RCL sequence has two domains that appear to have evolved separately from the MPP group (Supplementary Figure 6B). Notably, the two *Skoliomonas litria* M16 homologs branch with skoliomonad orthologs in two distinct clades. One of these branches from within a group of entirely bacterial PreP or MAG peptidase sequences and the other is a sister group of uncharacterized homologs from invertebrates (Metazoa) which lack predicted MTSs (except for one homolog in *Branchiostoma* that is the result of a recent duplication). This analysis suggests that none of the M16-related proteases in skoliomonads are orthologs of mitochondrial MPP or PreP; they are likely M16 proteases with separate cytosolic functions.

A candidate mitochondrial carrier protein (MCP) was identified in *Skoliomonas litria* but its substrate specificity could not be determined by phylogenetic analysis (Supplementary Figure 7). Whether this protein functions in the MRO membrane is unclear as MCPs are also known to function in peroxisomes or other subcellular organelles (Mazurek et al., 2010). Finally, a candidate homolog of Sam50 was identified in *Skoliomonas litria* and *Skoliomonas* sp. GEMRC. However, in phylogenetic analyses these proteins did not emerge within the clade of mitochondrial Sam50 orthologs, but instead formed an independent group emerging half-way between mitochondrial Sam50 and bacterial BamA clades (Supplementary Figure 8A). Orthologs of this Sam50/BamA-like protein were not detectable in any of the other BaSk genomes, even when this sequence was included in the HMM profile used for searching. To determine if this Sam50/BamA-like protein folded into a structure resembling the mitochondrial orthologs, we used AlphaFold 2.0 (Jumper et al., 2021) to predict its structure. The resulting structures (Supplementary Figure 8B, C) show that while these candidate Sam50 proteins possess what could be a intermembrane POTRA domain, the beta-barrels are smaller and adopt a different confirmation compared to Sam50 homologs of yeast (Supplementary Figure 9) and other well characterized examples (Takeda et al., 2021).

Among the five MRO predicted proteomes we present here, *Skoliomonas* sp. GEMRC and *Skoliomonas* sp. RCL contain the most candidate MRO translocon proteins, which is similar to the repertoire found in *Giardia intestinalis* (Leger et al., 2017) with the notable exception of the absence of GrpE in all members of BaSk. *Barthelona* sp. PCE, *Barthelona* sp. PAP020 and *Skoliomonas litria* possess a remarkably reduced MRO translocon complex by comparison. This may be due to the highly divergent nature of membrane bound MRO proteins, which makes detecting such proteins challenging (Pyrihová et al., 2018). Alternatively, in the case of *Skoliomonas litria* which lacks all the highly conserved GCS, ISC, protein import/refolding proteins of MROs, it is unclear if the Sam50/BamA-like protein, the MCP and MPP/M16 metalloprotease homologs are targeted to an organelle at all. It is possible that *Skoliomonas litria*, like the distantly related *Monocercomonoides exilis*, might completely lack a mitochondrial compartment and these proteins have taken on roles elsewhere in the cell. Further work needs to be undertaken to determine the cellular localization of these proteins and, if it exists, the function *Skoliomonas litria*’s MRO would carry out in the absence of all known conserved mitochondrial pathways and systems.

MTS prediction is a known challenge in inferring MRO function in anaerobic protists, including metamonads (Jerlström-Hultqvist et al., 2013; Schneider et al., 2011; Tanifuji et al., 2018), due to highly divergent or missing MTSs in these organisms. It has been suggested that the lack of a long positively charged MTS in metamonads is due to the loss of the proton gradient in the organelles (Garg et al., 2015), though recent analysis of canonical MTSs found in the Anaeramoebae – deep-branching metamonads – suggest that this may not always be the case (Stairs et al., 2021). Previous work has shown that some MRO proteins rely on cryptic internal signals to target proteins to the organelle with high fidelity (Garg et al., 2015; Mentel et al., 2008). Furthermore, most available MTS prediction software tools were trained on experimentally validated datasets of targeted proteins that are lacking for most protists. For these reasons, making accurate MTS predictions in metamonads is exceedingly difficult. Here, we employed several different MTS prediction software tools to find and annotate these signals in BaSks (see Supplementary Table 1). We found several examples of false positives in *Skoliomonas litria* and *Skoliomonas* sp. GEMRC, as homologs of transposons and nuclear targeting proteins were predicted to possess an MTS by some targeting prediction software with high confidence. Because of this, we considered proteins with MTS predictions from two or more MTS prediction software to be “strongly supported” targeting signals. Based on this criterion, *Skoliomonas litria* lacked any strongly supported targeting signals, while strongly supported targeting signals were rare in all other BaSk members. *Barthelona* sp. PCE contains strongly supported MTSs predicted on three proteins: an [FeFe]-hydrogenase, the T-protein of the glycine cleavage system (GCST), and on the IscU scaffold protein. *Barthelona* sp. PAP020 also contained a strongly supported MTS on its ortholog of IscU, as well as a different component of the glycine cleavage system (GCSP2). Another MTS detected was found on pre-sequence translocated-associated motor protein PAM18 in *Skoliomonas* sp. GEMRC. Interestingly, in *Skoliomonas* sp. RCL we find that a cytosolic CCT-gamma subunit contains a strongly predicted MTS, where in other BaSks, all cytosolic CCT subunits appear to be missing a MTS. However, when aligned to other BaSk CCT-gamma subunits, it does not appear to have an obvious N-terminal extension. As *Skoliomonas* sp. RCL has no mitochondrial Cpn60 homolog but does contain proteins which are homologous to MRO-localized proteins in other metamonads, it is possible (but unprecedented) that this CCT subunit has replaced the mitochondrial homolog in this species by acquiring an MTS. *Skoliomonas* sp. RCL does not have an additional copy of the gamma subunit of CCT. Follow up localization experiments are needed to deduce the subcellular localization of this particular CCT subunit.

### The BaSks share a core ATP generation pathway

All BaSk members share a similar hypothetical ATP production pathway that is reminiscent of the pathway found in *Giardia intestinalis* (Jedelský et al., 2011; Sanchez & Müller, 1996). In this pathway, pyruvate:ferredoxin oxidoreductase (PFO) catalyzes the oxidation of pyruvate in the presence of ferredoxin and CoA to produce acetyl-CoA, CO_2_ and reduced ferredoxin. Acetyl-CoA is then transformed to acetate by ACS and, in the process, converts ADP to ATP. Ferredoxin is reoxidized by [FeFe]-hydrogenase (HydA) that passes electrons to protons creating H_2_ gas. Notably, pyruvate-formate lyase (PFL) was also detected in all BaSk members, allowing an alternative pathway of acetyl-CoA production from pyruvate (Stairs et al., 2011; see Supplementary Figure 10). Note that there are multiple copies of both HydA and PFO in each BaSk member, some of which possess additional domains encoding ferredoxin or flavodoxin-like proteins (see Supplementary Table 2). No components of typical ATP production pathways found in trichomonad hydrogenosomes, such as acetate:succinyl-CoA transferase or succinyl-CoA synthase (Lahti et al., 1992; Van Grinsven et al., 2008), were detected in any of the BaSks. Though [FeFe]-hydrogenase is present in each BaSk member, none of the [FeFe]-hydrogenase maturase proteins (HydG, E or F) were detected. This is notable because these maturases, when present, are always targeted to mitochondria or MROs. This, combined with the lack of MTS detected in any of the proteins from the above pathway, leads us to assume that ATP and H_2_ gas are likely produced in the cytosol of the skoliomonads.

However, the pathways in the barthelonids appear to be slightly different in function and likely localization. Both *Barthelona* sp. PCE and *Barthelona* sp. PAP020 encode NuoE and NuoF NADH-dehydrogenase subunits of complex I that are known to function in the mitochondrial/MRO matrix in eukaryotes. It is widely thought that in the anaerobic protists that possess them, NuoE and NuoF likely form a complex with an MRO-localized HydA to perform simultaneous NADH and ferredoxin (Fd^-^) oxidation by an electron confurcating reaction that produces H_2_ gas, NAD+ and Fd (Dyall et al., 2004; Stairs et al., 2015). For this reason, and the fact that one of the HydA proteins of *Barthelona* sp. PCE and *Barthelona* sp. PAP020 have a predicted MTS, we infer that both barthelonids likely have MRO-associated hydrogenase activity, as recently suggested for *Barthelona* sp. PAP020 by Yazaki et al. (2019). However, unlike the hypothesis of Yazaki and colleagues (see Fig. 4b in Yazaki et al. 2019), we suggest that the reduced ferredoxin may, in fact, get produced within their MROs by PFO activity. If so, it is also possible that at least one of the two ACS homologs in each of these organisms is also MRO-localized and produces ATP within the organelle, though no MTSs are predicted on any of the ACS homologs. Further localization experiments in these barthelonids are needed to test these hypotheses.

**Figure 4.**
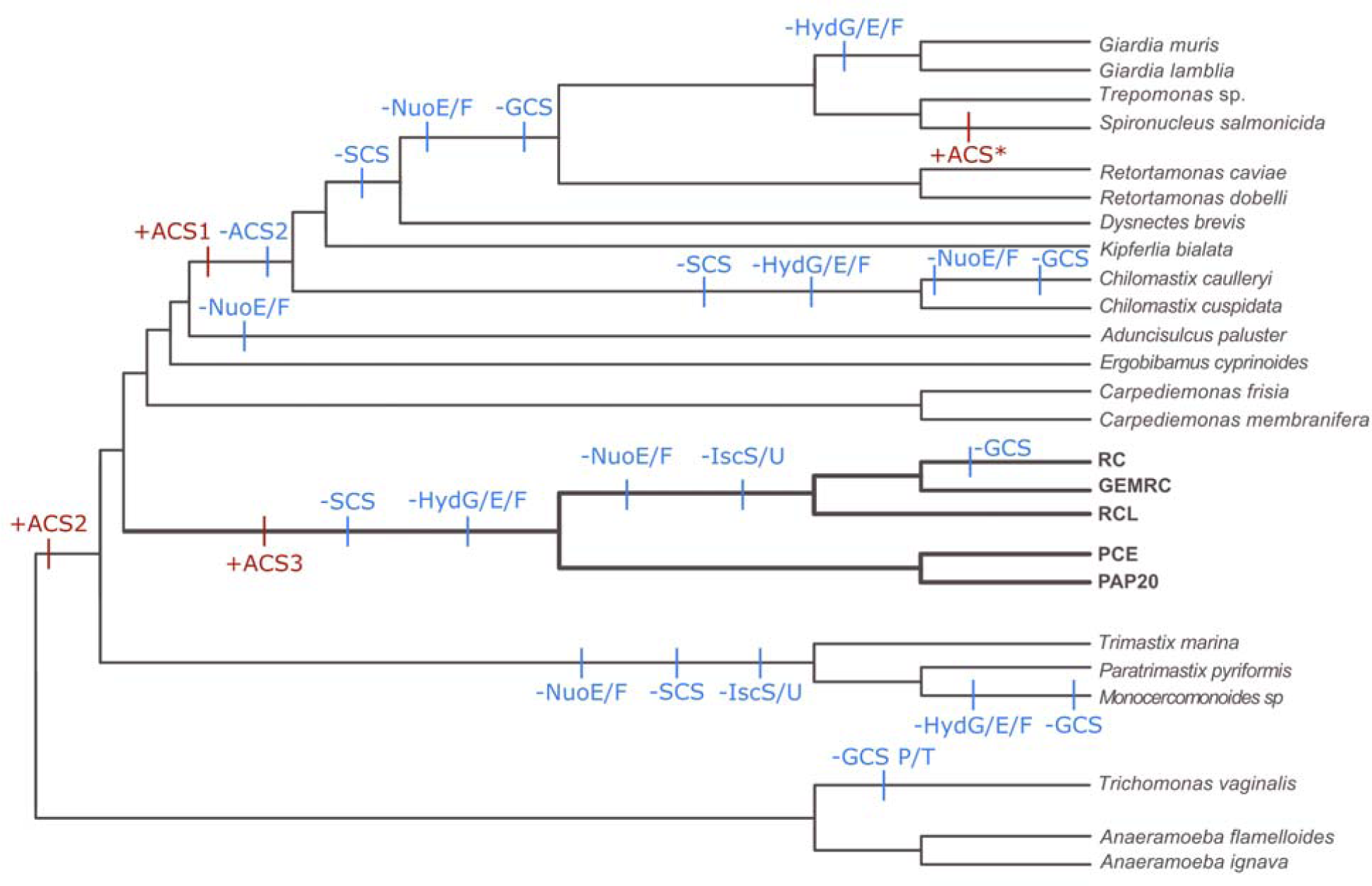
Gain and loss of anaerobic enzymes and MRO proteins over the tree of Metamonada. Gains (red) and losses (blue) of key metabolic pathway components discussed throughout this paper are mapped on a schematic phylogeny of Metamonada (based on Figure 1). Protein abbreviations are: HydG/E/F – Iron hydrogenase maturase G/E/F; ACS - Acetyl-CoA synthase; NuoE/F - Respiratory-chain NADH dehydrogenase 24/51 kDa subunit; GCS – Glycine cleavage system; SCS - Succinyl-CoA synthase;IscS/U - Iron-sulfur cluster assembly enzyme system. The Spironucleus ACS is denoted as ACS* as an ACS “type” could not be confidently assigned.

Previous studies have shown that some of the enzymes involved in ATP synthesis found within metamonads have a complex phylogenetic history. In particular, ACS is thought to have been transferred into the metamonads multiple times through LGT (Leger et al., 2017). To determine the evolutionary histories of the BaSk ACSs, we constructed a phylogenetic tree containing ACSs from a variety of metamonads, which can be found as Supplementary Figure 11. Our results show that, as in previous studies, most metamonad ACSs branch in one of two clades, known as ACS1 and ACS2. The ACS homologs found in the BaSks also form two distinct groups; one appears to be related to ACS2 found within *Carpediemonas membranifera*, *Ergobibamus cyprinoides,* and *Trimastix*. Additional ACS2 homologs from *Monocercomonoides* and *Paratrimastix pyriformis* were also identified, which also branch with the metamonad ACS2 group. This suggests that ACS2 was inherited in the last common ancestor of Preaxostyla and the fornicates + BaSk. The BaSk also possess a second ACS homolog which branches separately from other metamonad ACSs. This enzyme seems to be most closely related to ACS found within *Streblomastix*, *Monocercomonoides* and *Blastocystis*, and is distinct from other types of ACS in the tree. As previously reported by Yazaki et al. (2019) in their analysis of the *Barthelona* sp. PAP020 transcriptome, this is likely to be a novel homolog of ACS (ACS3) and was probably transferred into metamonads through a LGT event distinct from the event that gave rise to ACS1 and ACS2. Our analysis also suggests that the ACS found within *Spironucleus salmonicida* MROs may not be ACS2, as previously reported (Leger et al., 2017) because, in our analyses, the *S. salmonicida* ortholog (denoted ACS* in Figure 4 and Supplementary Figure 11) did not form a clade with other metamonad ACS2 sequences but instead grouped with several TACK archaea, although the support values for this relationship are low. A diagram summarizing the gains and losses of these proteins, along with other key metabolic processes and proteins mentioned above, can be found as Figure 4. These patterns suggest that there have been at least three, and possibly four, independent transfers of ACS into the metamonads. These transfers occurred at very different points in time along the evolution of various metamonad groups and highlights the role of LGT in the evolution of anaerobic metabolism.

To investigate the expression level of these key metabolic enzymes, we mapped RNASeq reads to each gene model, then normalized read counts and compared the fragments per kilobase of transcript per million mapped reads (FPKM) value amongst these gene models (see Supplementary Table 1). We found that some enzymes relating to anaerobic metabolism were amongst the top 100 most highly expressed genes in all four of the BaSk queried, though the specific anaerobic enzymes in this top 100 list differed between each species (see Supplementary Table 3). For *Skoliomonas litria*, *Skoliomonas* sp. GEMRC, and *Skoliomonas* sp. RCL this included two or more copies of genes encoding PFL. For *Skoliomonas litria* and *Skoliomonas* sp. RCL, this also included one copy each of a gene encoding PFO. *Skoliomonas litria*, in addition to PFL, had a copy of an ACS and a malic enzyme gene in its top 100 most highly expressed gene set. *Skoliomonas* sp. GEMRC was the only organism with a SmsCB gene represented its top 100 expressed list; other BaSk members have comparatively lower expression levels of this gene.

### Implications for the evolution of MROs within the Metamonada supergroup

Based on our reconstructions of MRO properties on the metamonad tree, we suggest that the last common ancestor of BaSk possessed an already highly reduced MRO compared to the common ancestor of fornicates + BaSk (Figure 4). The barthelonids PCE and PAP020 seem to have functionally reduced MROs, but their predicted MRO proteomes retain similar functions to other free-living metamonads such as the GCS and serine metabolism, hydrogen production and Fe/S cluster biogenesis by a simple ISC system. On the other hand, the MROs of *Skoliomonas* sp. RCL and *Skoliomonas* sp. GEMRC appear to be unique amongst metamonads, as we predict that the organelles only function in glycine and serine metabolism and are unlikely to produce H_2_. For *Skoliomonas litria*, the only proteins typically associated with mitochondria and MROs detected were a candidate MCP and a homolog of Sam50; of these, only the latter is exclusively associated with mitochondrial organelles in eukaryotes. The complete lack of all other proteins homologous to common MRO-localizing proteins could indicate that an MRO may not even exist in this organism. In any case, this extreme reductive evolutionary path may relate to the acquisition of a simple fused SmsCB gene that was acquired by LGT in a common ancestor of BaSk. This acquisition then led to the simplification of the ISC system in the *Barthelona* clade, and the outright loss of the ISC system in the skoliomonads, as Fe/S proteins for organellar functions were no longer needed.

Regardless of the evolutionary forces involved, the BaSks demonstrate reduction in MRO function that is independent of the loss seen in other metamonad groups, such as diplomonads and oxymonads (Karnkowska et al., 2016; Leger et al., 2017). This is remarkable given that many metamonads that display a similar loss of MRO functions are parasitic or commensalistic and the nutrient-rich endobiotic lifestyle has often been assumed to be part of the explanation for extreme MRO proteome streamlining (Karnkowska et al., 2016; Stairs et al., 2015). Instead, in the case of the free-living BaSk members, it appears that gain of a simple SMS system was amongst the first triggers leading to the extreme reductive MRO evolution in this group.

## Conclusions

We have generated high quality draft genomes, transcriptomes, and curated predicted proteome sets for a novel group of free-living metamonads. The placement of BaSk within the tree of Metamonada as the sister group to the Fornicata make them a key group to study in terms of genome evolution, LGT events, and adaptation to various environmental niches. These data place us several steps forward towards the goal of representing diverse and historically under-sampled microbial eukaryotes in the tree of eukaryotic life.

Our findings demonstrate the diversity of MRO configuration and function amongst the metamonads. As a group, BaSks display a level of MRO reduction not typically seen in free-living metamonads and which is independent of the reduction displayed in the parasitic diplomonads. BaSk members share many interesting parallels with the MRO reduction and loss seen in other metamonads, as all BaSks possess a SMS Fe/S cluster system that has, in some cases, replaced the canonical ISC system. This is the second discovered instance of an LGT event that has replaced the ISC system in metamonads with a SUF-like system and serves to highlight the role of adaptive LGTs in MRO evolution. Based on these findings, we speculate that the last common ancestor of the BaSks possessed both a compact, streamlined genome with a highly reduced MRO, and was markedly different from the last common ancestor of the fornicates. In many of the BaSk taxa, further work is needed to elucidate the exact subcellular localization of several candidate MRO proteins including enzymes in the anaerobic ATP synthesis pathway and to determine the breadth of functions carried out by the MROs in this newly discovered group. For *Skoliomonas litria*, which lacks all known mitochondrial pathways, it will be especially important to investigate the localization of its anaerobic ATP synthesis pathway, its MCP, and the Sam50/BamA-like protein to determine whether this free-living flagellate has completely dispensed with the organelle.

## Materials and Methods

### Cell cultivation & microscopy

Established cultures of isolates *Skoliomonas litria*, *Skoliomonas* sp. RCL and *Skoliomonas* sp. GEMRC (Eglit et al. 2024) were maintained in variants of CR media (Gigeroff et al., 2023); *Skoliomonas litria* in 50 mL of 40 ppt CR with 3% LB, *Skoliomonas* sp. GEMRC in 50 mL of 40 ppt CR with 1% LB and 3 sterile wheat grains, *Skoliomonas* sp. RCL in 50 mL of 25 ppt CR with 1% LB and 3 sterile wheat grains. All were inoculated into fresh media every 7-10 days based on culture density, with a second inoculation after 24 hours. Cultures of isolate ‘*Barthelona* sp.’ PCE (Yazaki et al. 2019) were maintained in sterile filtered seawater (Northwest Arm, Halifax, Nova Scotia) supplemented with 3% LB, and kept at 16L. Cell growth was monitored weekly, and subculturing was done every 3-4 weeks depending on cell density by transferring around 20% of the old culture into the new culture.

Cells were imaged with differential interference contrast optics on a Zeiss Axiovert 200M microscope fitted with a AxioCam M5 camera (Carl Zeiss AG).

### Nucleic acid extraction & sequencing

For *Skoliomonas litria*, *Skoliomonas* sp. GEMRC, *Skoliomonas* sp. RCL, and *Barthelona* sp. PCE: cell material for DNA and RNA extraction was harvested from 1-2 L of dense cell culture. The cells were collected by centrifugation in 50 mL tubes for 8 min at 2000×*g* and at 4L. The pellets were then combined into two 15 mL tubes that were spun again as before. The recovered pellets were resuspended in sterile filtered (0.2 µm) spent culture media and layered over Histopaque-1077, and then subjected to centrifugation at 2000×*g* for 20 min at room temperature. The protist-containing layer between the Histopaque and the top media was collected, and the procedure was repeated for a total of two rounds of Histopaque separation. Finally, the cells were diluted by 5 mL (10 volumes) of sterile spent-media and pelleted for 8 min, at 2000×*g*, 4L. The cells were then resuspended in 2 mL of fresh media.

Genomic DNA was purified using a variety of techniques including traditional phenol-chloroform extraction, CTAB extraction, the QIAGEN Genomic-tip kit, and the QIAGEN MagAttract HMW DNA kit. RNA was extracted using Trizol according to Invitrogen specifications. RNA and DNA were sent to Genome Quebec for library construction and NovaSeq 6000 Illumina sequencing. In the case of the RNA, poly-A selection was performed to enrich for eukaryotic reads. High molecular weight DNA from each species was used to construct a 1D ligation Nanopore library (LSK108 for *Barthelona* sp. PCE, LSK109 - EXP- NBD104 for *Skoliomonas litria* and *Skoliomonas* sp. GEMRC, and LSK308 for *Skoliomonas* sp. RCL) and was sequenced using a MinION flowcell (FLO-MIN106 (R9.4) for *Skoliomonas litria* and *Skoliomonas* sp. GEMRC, and FLO-MIN107 (R9.5) for *Skoliomonas* sp. RCL and *Barthelona* sp. PCE).

For *Barthelona* sp. PAP020, cells were cultured as described in Yazaki et al. (2019). Cells were harvested by pelleting at 1500 rpm for 5 minutes at 4L and were washed with PBS prior to phenol chloroform DNA extraction. The purified DNA was sequenced on an Illumina Hiseq 2500 PE using the Trueseq library prep.

### Genome/transcriptome assembly, & gene annotation

For *Skoliomonas litria*, *Skoliomonas* sp. GEMRC, *Skoliomonas* sp. RCL, and *Barthelona* sp. PCE, Illumina NovaSeq reads were trimmed using Trimmomatic v0.36 (Bolger et al., 2014). RNASeq reads were assembled using Trinity v2.6.6 (Haas et al., 2013) and checked for multiplexing contamination using WinstonCleaner (https://github.com/kolecko007/WinstonCleaner). Bacterial contamination was removed from the Trinity assemblies using Anvi’o v5 (Eren et al., 2021).

Raw MinION sequencing data was basecalled using Albacore v2.1.3 (Albacore has since been replaced by Guppy – see Wick et al., 2019). These data were assembled using Flye v2.3 (Kolmogorov et al., 2019), Raven v0 (Vaser & Šikić, 2021) Canu v1.7 (Koren et al., 2017) and ABruijn v2.2b (Lin et al., 2016). The resulting assemblies were evaluated for completeness and contiguity by comparing assembly size, contig number, N50 and L50 values, and ALE scores (Clark et al., 2013). The ABruijn assembly was chosen moving forward for *Skoliomonas litria*, *Skoliomonas* sp. RCL and *Barthelona* sp. PCE. For *Skoliomonas* sp. GEMRC, the Canu assembly was used. Raw MinION output was used to improve the base-call accuracy of these assemblies using Nanopolish version 0.8.4 (Loman et al., 2015) and Illumina reads were used to error-correct the assemblies using Unicycler version 0.4.3 (Wick et al., 2016). Bacterial contamination was assessed and removed from the resulting contigs using read coverage, GC content, and BLAST search results (Altschul et al., 1990). The Eukfinder workflow was used to remove prokaryotic contaminants from the BaSk genome assemblies (Zhao et al., 2023)

Gene prediction utilized a pipeline described in Salas-Leiva et al. (2021). Briefly, RepeatMasker (Flynn et al., 2020) was first used to mask the repetitive regions of the genome. RNA-seq reads were then mapped to the genome using HISAT2 (Kim et al., 2019). GeneMark-ET (Lomsadze et al., 2014) then utilized this mapping information to generate gene coordinates and intron splice sites, creating a training set of genes. This training set was then used to train AUGUSTUS (Stanke et al., 2004), which generated the gene predictions. PASA (Haas et al., 2003) was then used to validate the coordinates of these gene models. The predicted protein set was checked for completeness using the obd9 BUSCO v3.0.1 database (Simao et al., 2015).

To estimate genome completeness we used Merqury v.1.0 (Rhie et al., 2020) with a kmer size of 17 (estimated using Merqury). The Illumina reads used for the Merqury analysis were decontaminated using the Eukfinder workflow (Zhao et al., 2023) before genome completeness estimation. Additionally, RNA-seq mapping was used as a measure of genome completeness, and RNA-seq reads were decontaminated using the same method mentioned used for the DNA Illumina reads.

To construct the *Barthelona* sp. PAP020 genome, Illumina reads were assembled using Spades (Prjibelski et al., 2020) with default parameters. Bacterial reads were identified by metaBAT (Kang et al., 2015) and CheckM (Parks et al., 2015). Once the identified bacterial reads were removed, the remaining Illumina reads were then used to assemble a new genome. This process was repeated until no bacterial contamination was detected. The final contigs were verified by checking kmer content and were also checked against the *Barthelona* sp. PAP020 transcriptome (Yazaki et al., 2020).

### Gene expression analysis

Expression of predicted protein-coding genes was quantified by mapping RNASeq reads to their respective long-read genome using HISAT2 (Kim et al., 2019). Using this mapping information and gene model coordinates, the expression level was normalized using Cufflinks (Cuffquant and Cuffnorm with default settings)(Trapnell et al., 2010) to compute a FPKM value for each gene.

### Phylogenomic analysis

The phylogenomic dataset was constructed based on a previously published set of 351 highly conserved protein orthologs (Brown et al., 2018), with 177 orthologs excluded due to deep paralogies or insufficient coverage across pertinent species in the analysis. Each of the 174 remaining ortholog alignments were used to construct single-protein phylogenies, and were used to screen out non-orthologous data. The final 174 alignments were realigned using mafft v7.310 einsi, trimmed using BMGE v1.0 (default settings) (Criscuolo & Gribaldo, 2010), and were concatenated together and used to infer a maximum-likelihood tree under the LG+C60+F+gamma model of evolution with IQTree v1.5.5 (Nguyen et al., 2015). A PMSF model based on this tree and mixture model (Wang et al., 2018) was estimated and used to calculate branch supports using non-parametric bootstrapping, approximate Bayes (aBayes; Anisimova et al., 2011) and approximate likelihood-ratio support (aLRT; Guindon et al., 2010).

### Inventory of MRO and anaerobic metabolism proteins

Proteins of interest were searched for in the predicted proteome, transcriptome, and genome of each species using both NCBI BLAST (using blastp, tblastn and blastx) and HMM profile searching using Hmmer v3.1b2 (hmmer.org). For each search, queries included both orthologous sequences from closely related metamonads and, once identified by these methods, the orthologs from other BaSks. The protein sets of the BaSk taxa were also used as queries in BLAST searches against the mitochondrial proteomes from *Andalucia godoyi* (Genbank accession VRVR00000000; see Gray et al., 2019), *Tetrahymena thermophila* (predicted protein Genbank accession EAR80512 to EAS07932, see Eisen et al., 2006; Smith et al., 2007), *Arabidopsis thaliana* (NCBI Bioproject PRJNA10719 and SAMN03081427; see Cheng et al., 2017; The Arabidopsis Genome Iniative, 2000), *Homo sapiens* (International Human Genome Sequencing Consortium, 2001; mitochondrial proteins from UniProt (GO:0005739)), *Acanthamoeba castellanii* (NCBI Bioproject PRJNA599339 and PRJNA487265; see Gawryluk et al., 2014; Matthey-Doret et al., 2022), and *Saccharomyces cerevisiae* (https://www.yeastgenome.org/; see Engel et al., 2014). The top five hits for each query sequence were examined manually. Proteins relating to mitochondrial protein translocation were searched for using both general HMM profiles (Leger et al., 2017) and profiles tailored for searching in metamonads. The latter profiles were created by retrieving the profiles from Pfam for each protein of interest and then retrieving the best hits using the Pfam profile against the several publicly available metamonad sequencing data databases with hmmsearch. Using an evalue cutoff of <0.01, metamonad sequences were added to the corresponding Pfam seed alignment using the mafft –add function. If there were more than two detected sequences from metamonads, an alignment was made with these sequences and was then used to build a profile to search for more hits. Each candidate protein identified was used to reciprocally search the BLAST database to confirm protein identity. In ambiguous cases, the Pfam and InterPro databases were searched to analyze protein domain structure.

Phylogenies for key proteins identified were constructed to aid in identification in cases with deep paralogy (Hsp70 and Cpn60). Phylogenies were also made to identify the origin of enzymes that act in anaerobic metabolism. Alignments from Leger et al. 2017 or Tsaousis et al., 2014 were utilized as a starting point, and we added the corresponding BaSk sequences. Where relevant, the protein sequences of the top 10 BLAST hits from each BaSk homolog were also added to the appropriate alignments. In cases where no starting alignments were available, a preliminary alignment was created by retrieving the top 1000 blast hits and reducing the sequences to 100-120 sequences using CD-HIT v.4.6 with recommended word sizes. These sequences were aligned using mafft einsi with default settings and were trimmed using trimal v1.4.rev15 (Capella-Gutiérrez et al., 2009) with the -gappyout setting. These phylogenies were inferred using the LG+C20+F+gamma model of evolution with IQ-TREE.

### Mitochondrion-related organelle protein localization predictions

Putative mitochondrial targeted proteins and mitochondrial targeting sequences (MTS) were predicted using the following mitochondrial targeting peptide prediction programs: TargetP v1.1b (Emanuelsson et al., 2007), MitoProt II v1.101 (Claros & Vincens, 1996), and MitoFates v1.1 (Fukasawa et al., 2015)). Additionally, the predicted proteins were used to search both the nr database and a custom database of known mitochondrial proteins.

### Protein structure prediction and alignment

The structure of candidate Sam50/BamA-like proteins in *Skoliomonas litria* and *Skoliomonas* sp. GEMRC were predicted using the ColabFold v1.5.2 server (Mirdita et al., 2022). The β-barrel portion of the *Skoliomonas litria* Sam50/BamA-like protein was aligned to the yeast Sam50 protein structure (Takeda et al., 2021) using the PyMOL “cealign” function.

### SMSCB (SUFCB) phylogeny

The SmsCB fusion protein identified in each BaSk member was added to previously constructed alignments (Leger et al., 2016) of SmsC and SmsBD. This dataset was aligned using mafft einsi with default settings and was trimmed by hand. The dataset was then concatenated together, and the phylogeny was inferred using the same methods as the single protein phylogenies mentioned above.

### M16 protease phylogeny

The MPP candidate proteins identified in the skoliomonads were added to the previously constructed alignments of M16 proteases (Garrido et al., 2022), which include SPP, PreP, MPP-alpha and MPP-beta sequences. In addition, we used the protein sequences of the two MPP candidates of *Skoliomonas litria* to BLAST search nr, and added the top 20 hits for each candidate to the alignments. This dataset was aligned using mafft einsi with default settings and was trimmed by trimal as above. After trimming, we removed the duplicated protease domains in four domain proteins, relative to their position when aligned to two domain proteins. This alignment was then inferred using the same methods as the single protein phylogenies described above. The domain structures displayed to the right of the phylogeny represents the predicted protein domains of the skoliomonads (in red) and other major clades within the tree (in grey) as predicted by the InterProScan online tool (see https://www.ebi.ac.uk/interpro/), which utilizes the InterPro 98.0 database (Paysan-Lafosse et al., 2023).

## Data availability

Genomes and gene annotations (where applicable) can be found under NBCI BioProject ID PRJNA949547.

## Acknowledgments

The majority of work was supported by Foundation grant (FRN-142349) from the Canadian Institutes for Health Research (CIHR) awarded to Andrew J. Roger. Shelby K. Williams was supported by a graduate scholarship from the Killam Foundation.

Supplementary tables 1-3: Available for download from Dryad https://datadryad.org/stash/share/SNhoYuY19An041SeN194bwM_K3WIqqOO4TwltqK1mn0

## Supplementary figures

**Supp. Figure 1.**
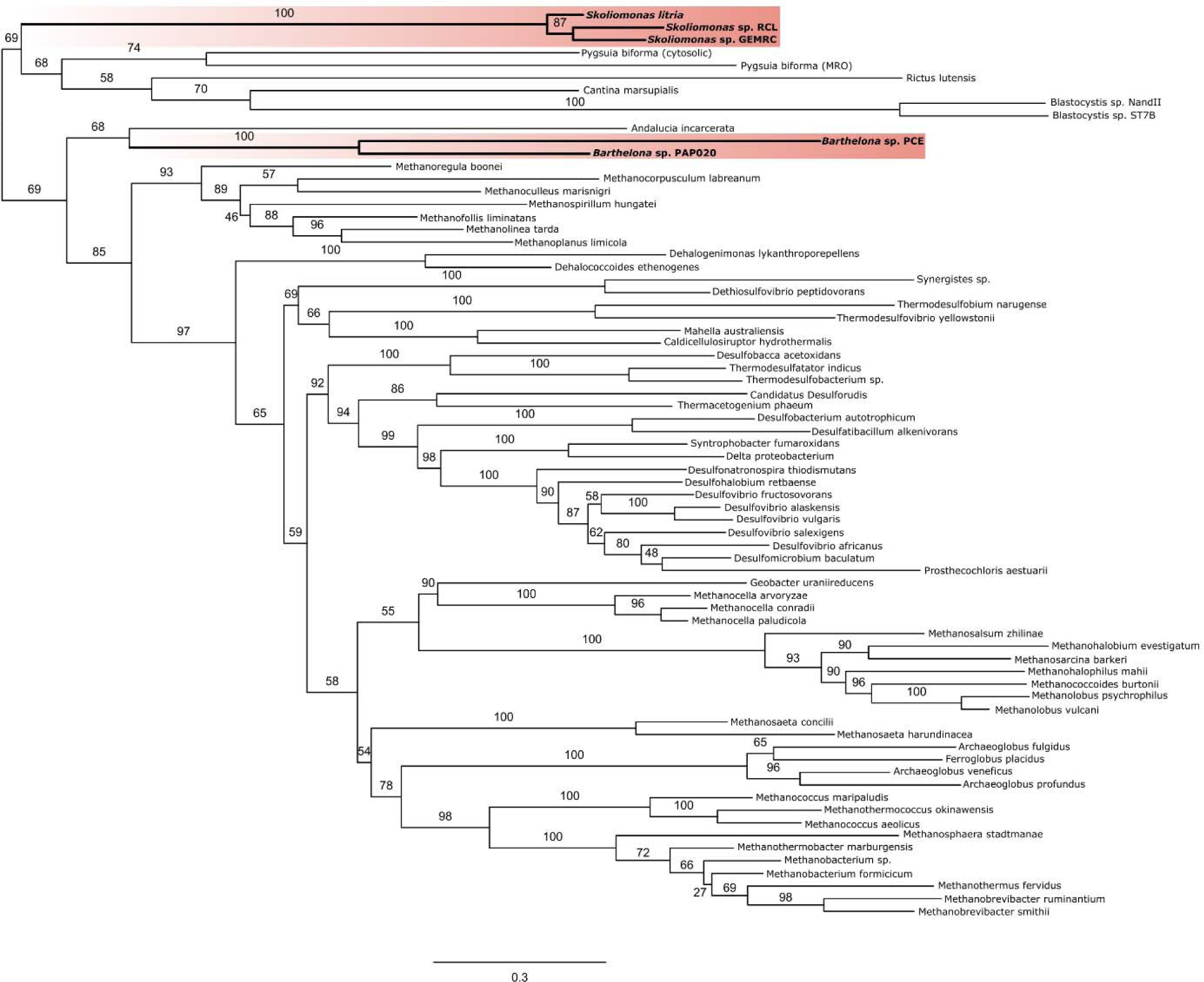
A phylogeny of the SmsC protein from eukaryotes, bacteria, and archaea. The ML tree was estimated using IQ-TREE under the LG+C60+F+Γ model of evolution. Ultrafast bootstrap values are displayed on the branches. For eukaryotic homologs, the N-terminal portion of the protein of the eukaryote SmsCB fusion that aligned with prokaryotic SmsC was used in the alignment. BaSk sequences are bolded and highlighted in red.

**Supp. Figure 2.**
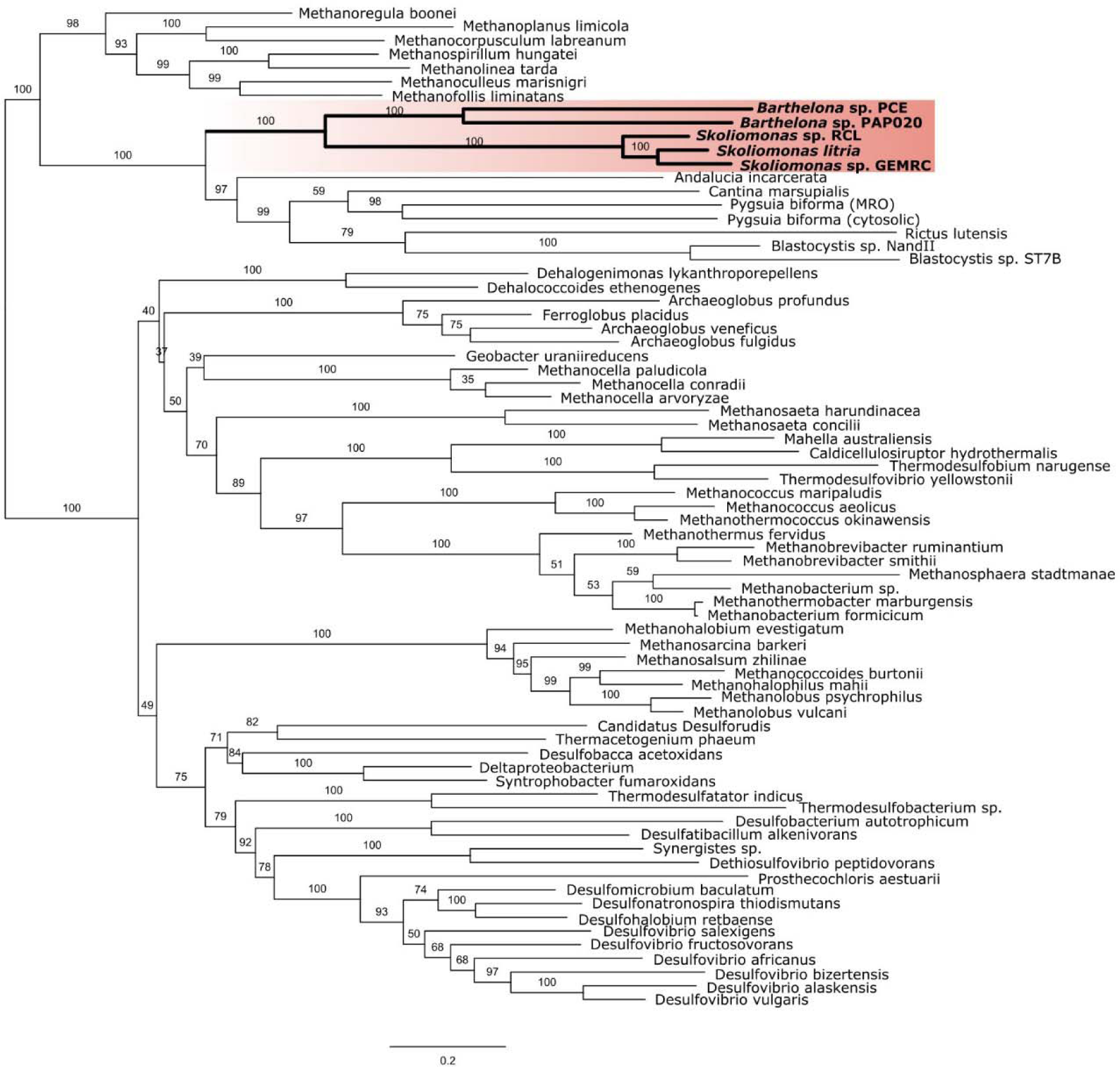
A phylogeny of SmsBD protein from eukaryotes, bacteria, and archaea, The ML tree was estimated using IQ-TREE under the LG+C20+F+Γ model of evolution. Ultrafast bootstrap values are displayed on the branches. For eukaryotic homologs, the C-terminal portion of the protein of the eukaryote SmsCB fusion that aligned with prokaryotic SmsB was used in the alignment. BaSk sequences are bolded and highlighted in red.

**Supp. Figure 3.**
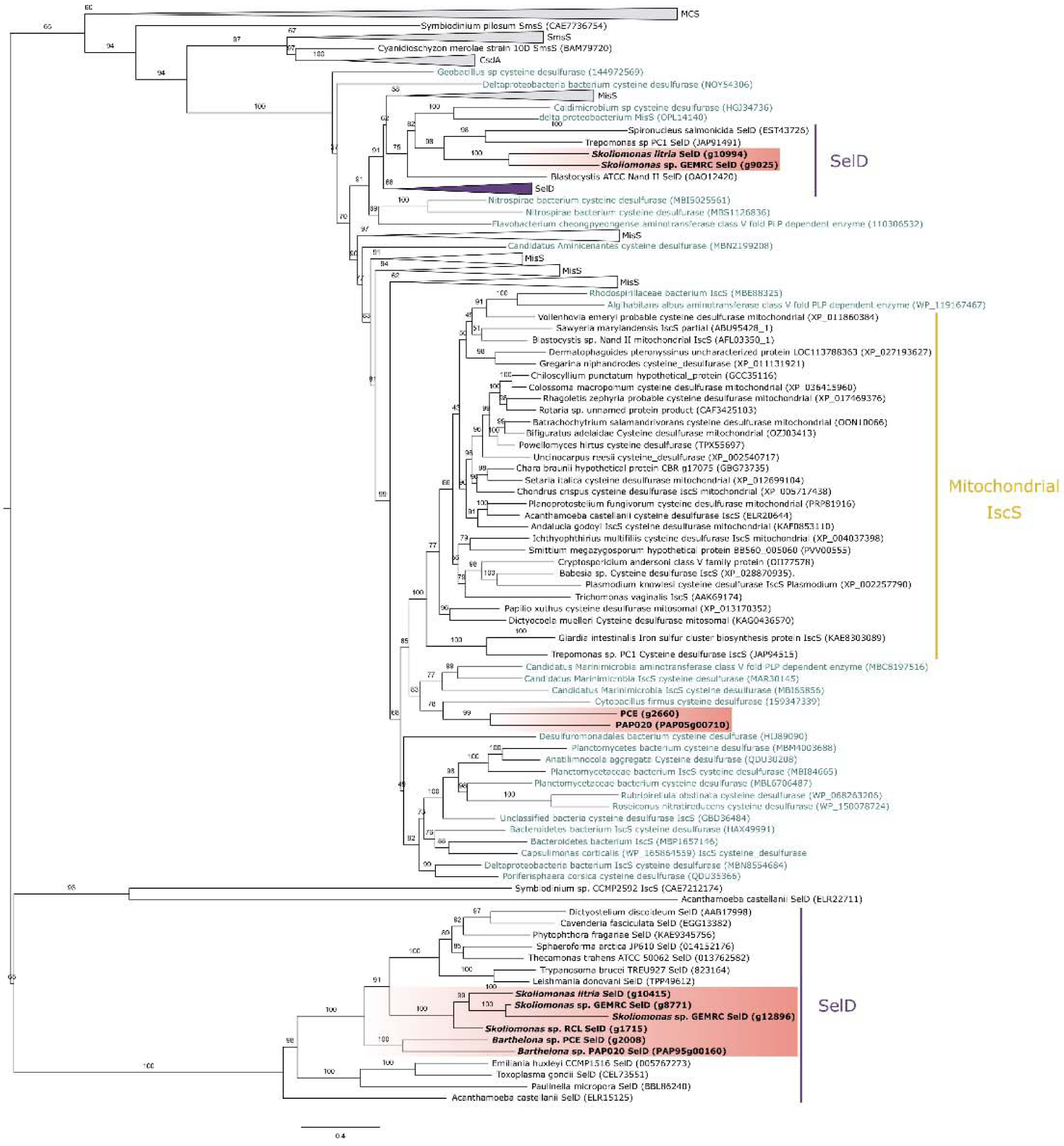
Phylogenetic relationships amongst several different types of cysteine desulfurase enzymes and several enzymes that share structural similarities with them. The ML tree was estimated using IQ-TREE with the LG+C20+F+Γ evolutionary model. Ultrafast bootstrap values are displayed on the branches. Several families of related proteins have been collapsed into wedges. BaSk sequences are bolded and highlighted in red. Abbreviations are: MCS – Molybdenum cofactor sulfurase; SmsS – SUF-like minimal system; CsdA - Cysteine sulfinate desulfinase A; MisS - Minimal iron-sulfur system; IscS - Iron-sulfur cluster assembly enzyme; SelD – Selenide water dikinase

**Supp. Figure 4.**
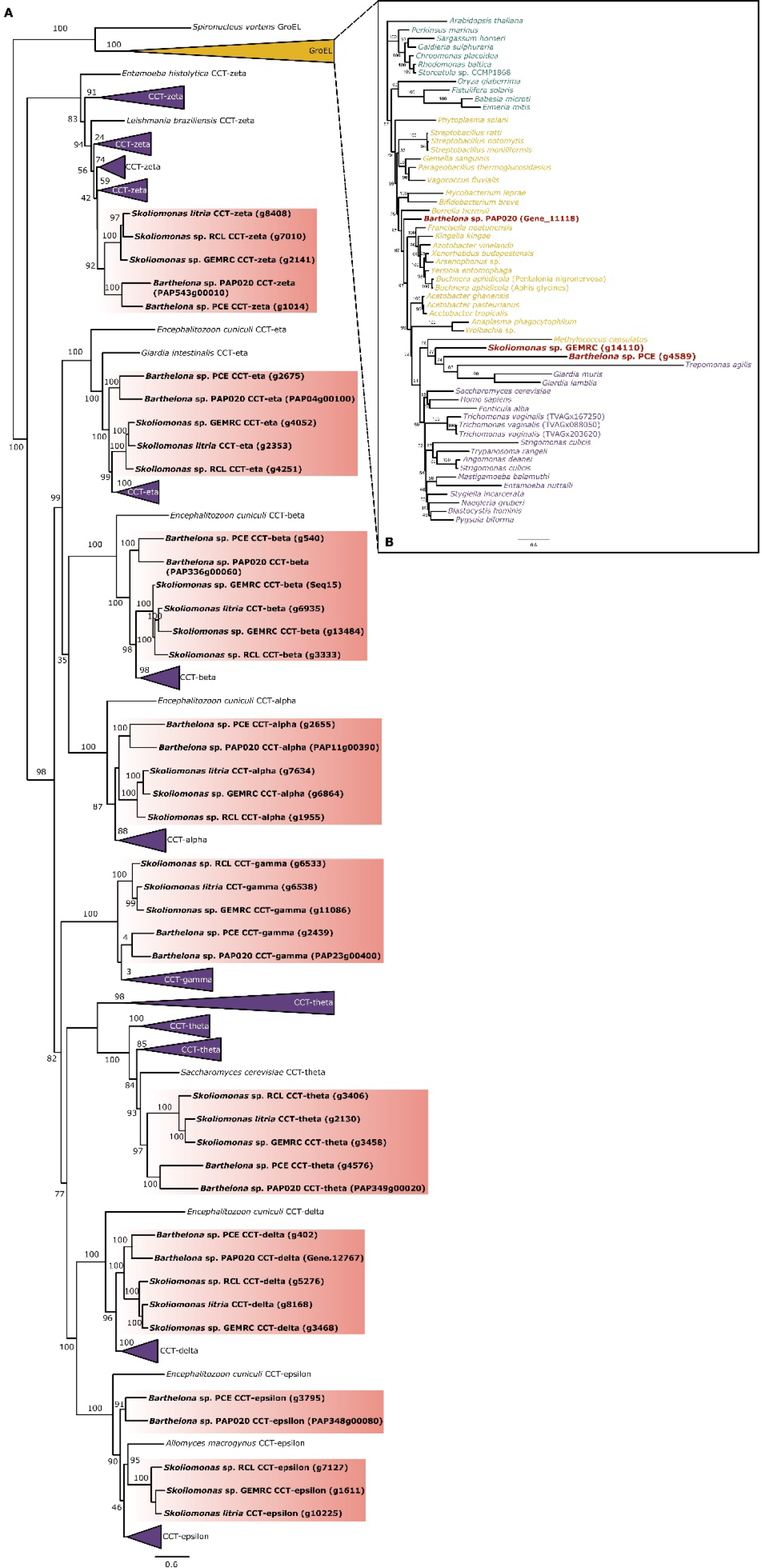
Phylogenetic relationships amongst chaperonin 60 (cpn60) homologs in eukaryotes. These ML trees were estimated using IQ-TREE with the LG+C20+F+Γ evolutionary model. Ultrafast bootstrap values are displayed on the branches. BaSk sequences are bolded and highlighted in red. (A). Both cytosolic (purple) and mitochondrial (orange) homologs are included in this tree. The major groups of proteins are displayed as wedges. (B). A reconstruction of the mitochondrial (purple), bacterial (orange), and choroplasatic (green) Cpn60 subunit GroEL.

**Supp. Figure 5.**
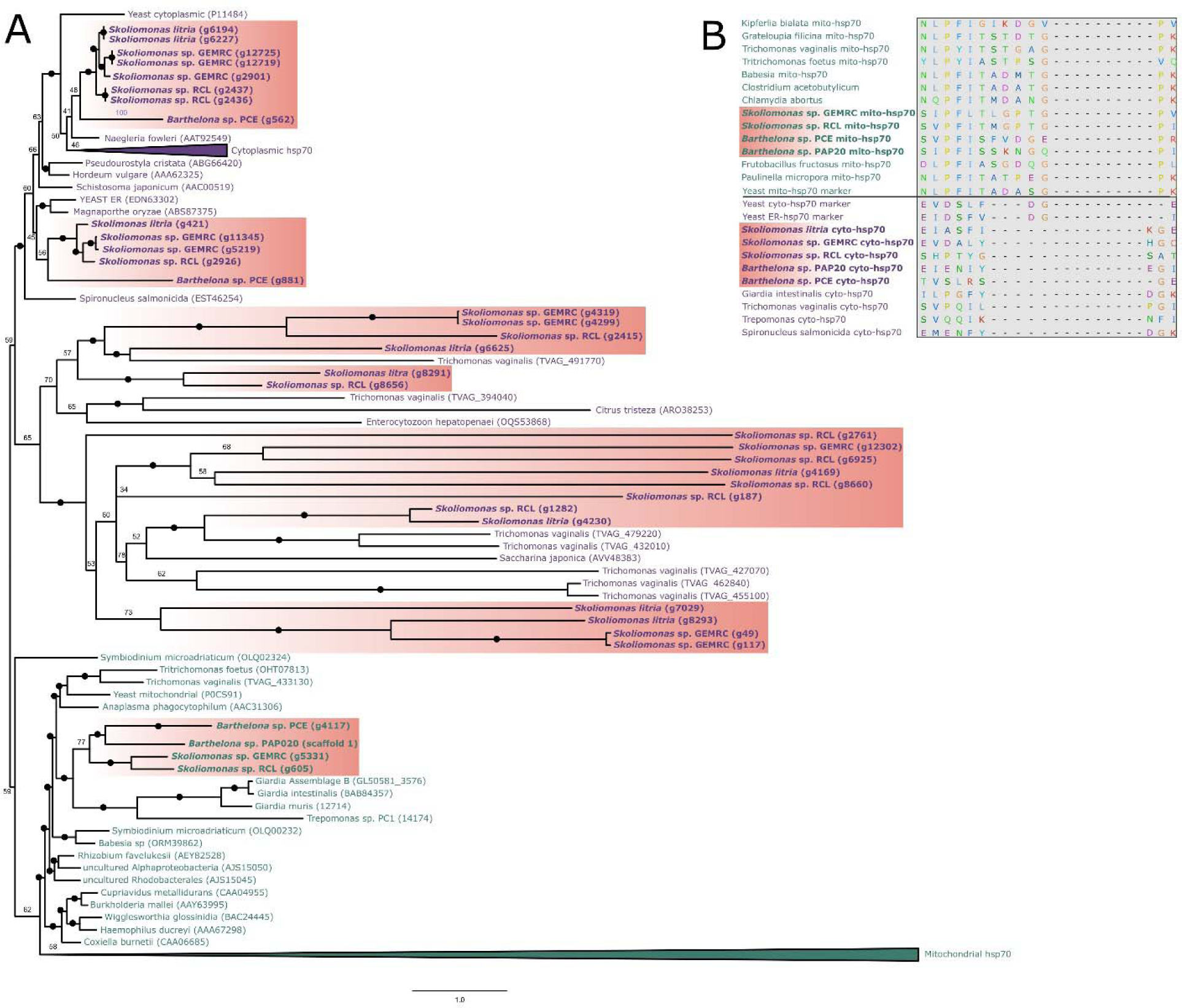
Phylogenetic relationships amongst heat shock protein 70 (hsp70) paralogs in eukaryotes and prokaryotes. (A) The ML tree was estimated using IQ-TREE with the LG+C20+F+Γ evolutionary model. Ultrafast bootstrap values are displayed on the branches. Cytoplasmic and ER (purple) and mitochondrial (green) homologs are labeled according to indel mapping patterns in the alignment. BaSk sequences are bolded and highlighted in red. (B) An example of indel patterns within the alignment used to assign probable localizations of hsp70 homologs in the tree. Amino acid residues are colour coded.

**Supp. Figure 6.**
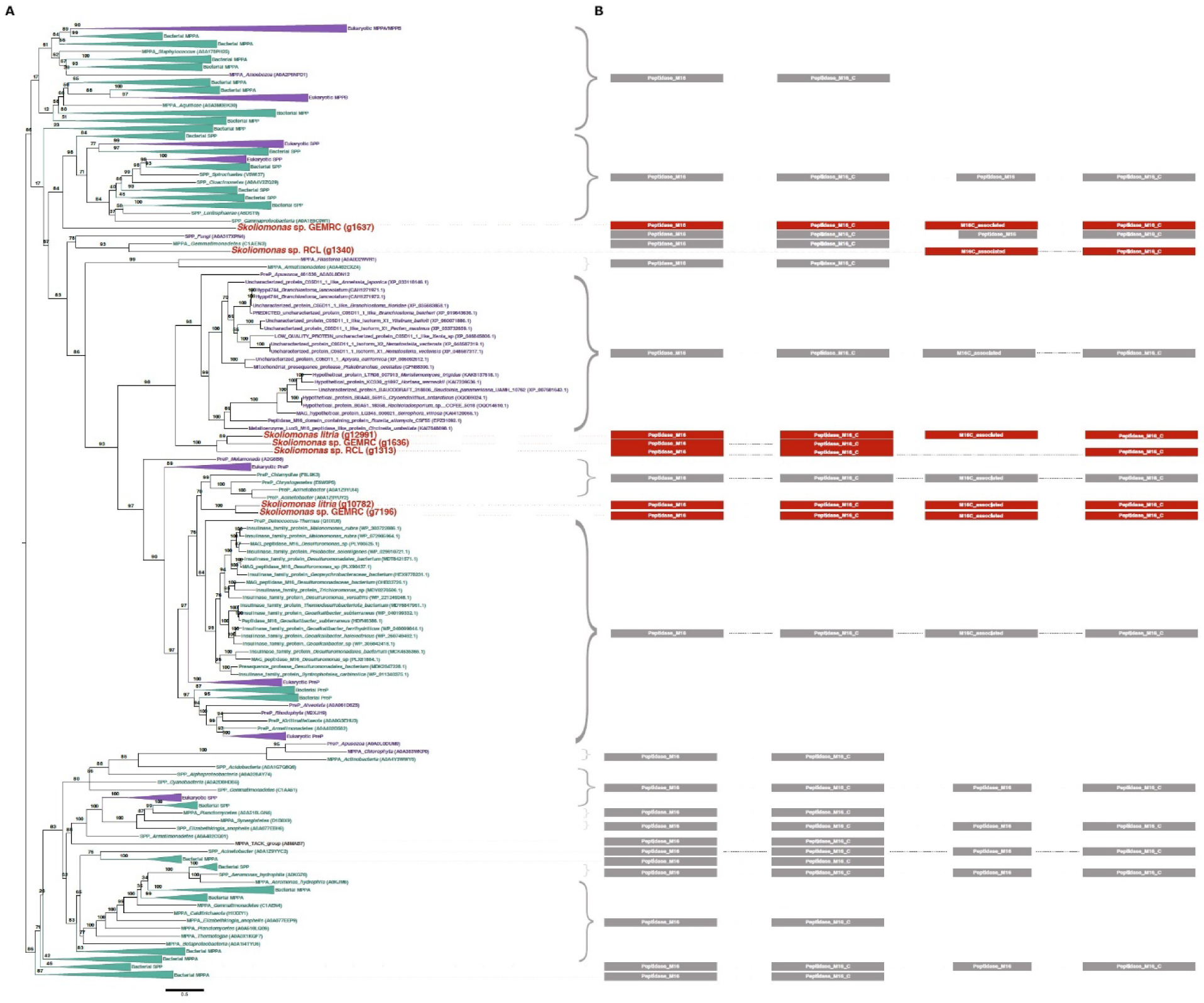
Estimated phylogenetic relationships amongst a variety of M16 metallopeptidases. This tree was estimated using the M16 peptidase database previously constructed by Garrido and colleagues (2022) and was estimated using IQ-TREE under the LG+C20+F+Γ evolutionary model. Ultrafast bootstrap values are displayed on the branches. Major groups of proteins have been collapsed into wedges. To the right of the tree, the domain structure of each protein/protein group is displayed, as annotated by InterPro. Eukaryotes – purple; bacteria – green; skoliomonads – red.

**Supp. Figure 7.**
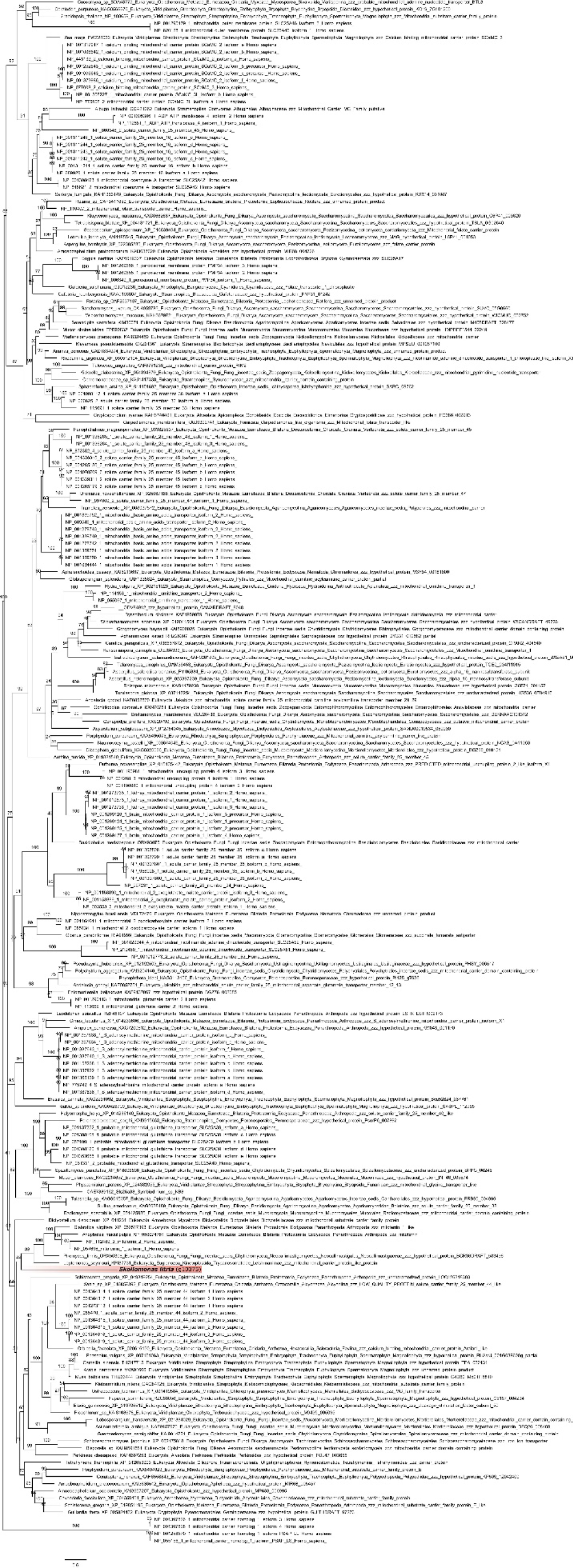
Estimated phylogenetic relationships between members of the mitochondrial carrier family (MCP) with candidate MCP sequences from ‘BaSks’. Where possible, MCP sequences are labeled with their family subtype designation according to the NCBI nr database. This phylogenetic tree was constructed using IQ-TREE and the LG+C20+F+Γ model of evolution. The *Skoliomonas litria* candidate MCP is highlighted in red. Values on the branches represent ultrafast bootstrap values.

**Supp. Figure 8.**
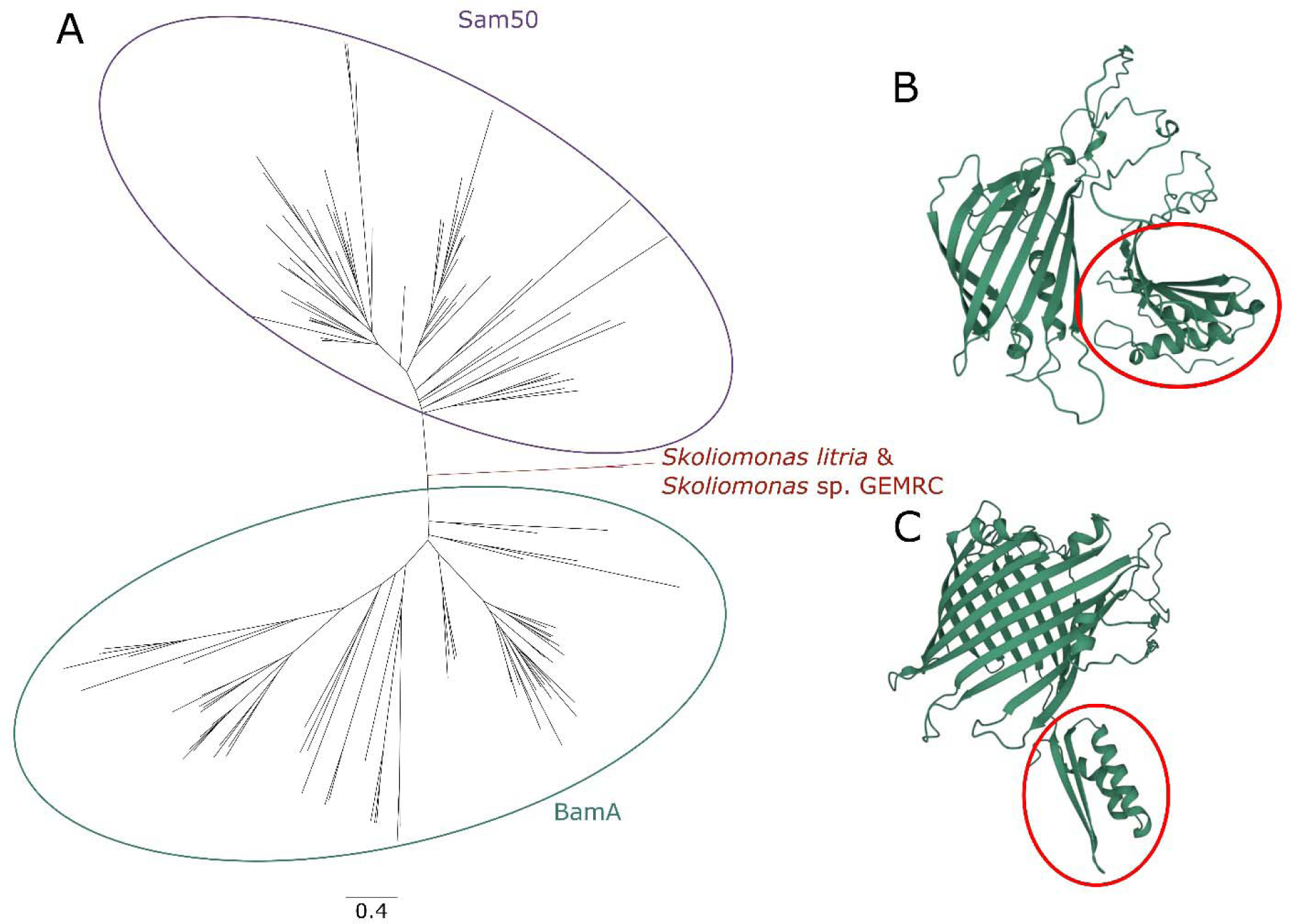
The phylogeny of Sam50/BamA family in eukaryotes and prokaryotes and predicted structures of the BaSk homolog. (A) A phylogenetic reconstruction of the Sam50 candidate proteins in *Skoliomonas litria* and *Skoliomonas* sp. GEMRC, and their relationship to Sam50 (purple) and BamA (green) homolog proteins in eukaryotes and prokaryotes. (B) Alphafold2 structure prediction using the sequence of the candidate Sam50 protein found in *Skoliomonas litria* (C) Alphafold2 structure prediction using the sequence of the candidate Sam50 protein found in *Skoliomonas* sp. GEMRC. The protein regions resembling “POTRA” domains in other Sam50 proteins are highlighted with a red circle in each candidate.

**Supp. Figure 9.**
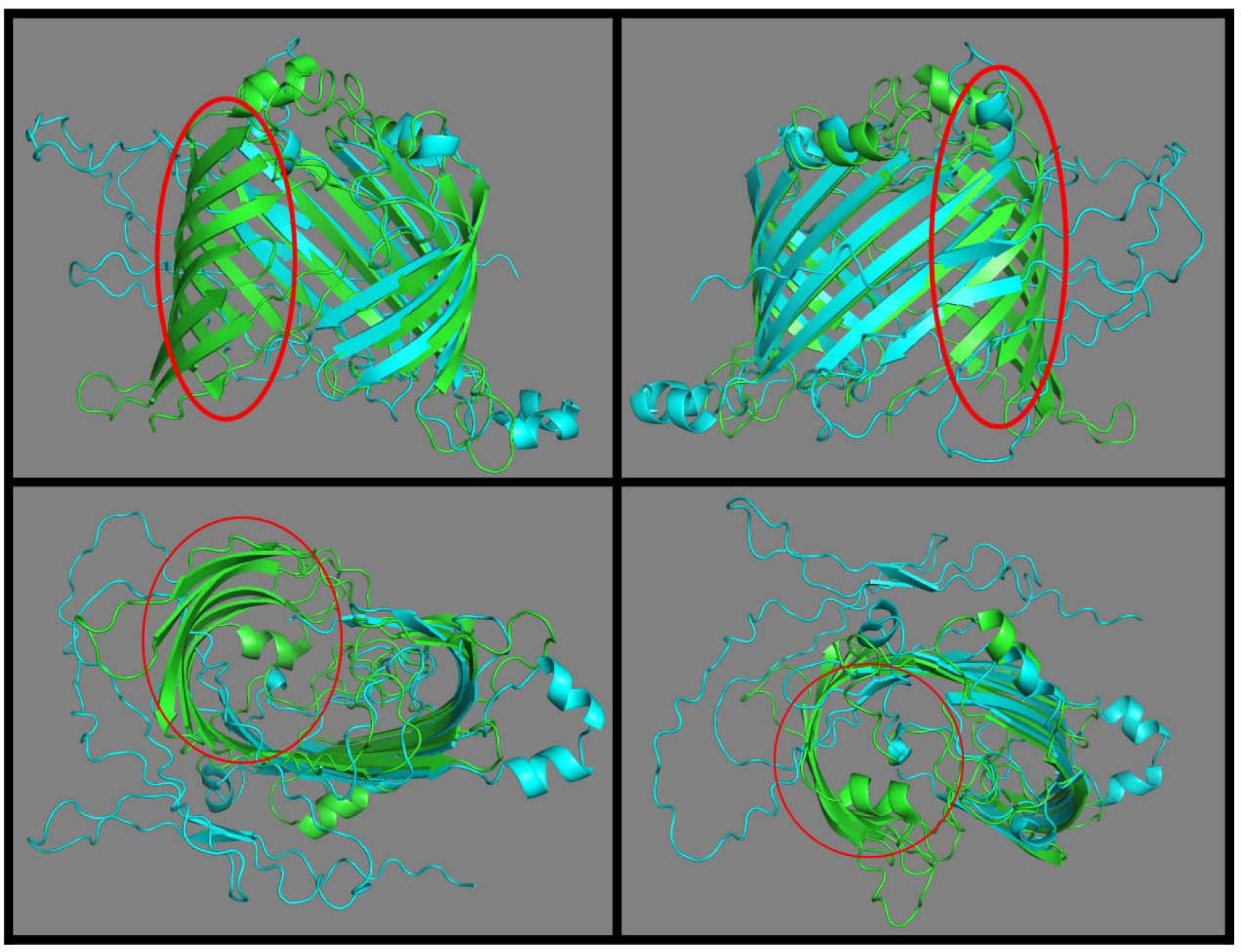
A structural alignment of the. β**-barrel portions of the *Skoliomonas litria* candidate Sam50 protein and Sam50 from yeast.** The *Skoliomonas litria* protein is shown in blue, aligned to the β-barrel domain of the yeast Sam50, which is shown in green. The missing sections of the β-barrel in *Skoliomonas litria*, when compared to the yeast model, is highlighted by a red circle. This structural alignment was produced using the PyMOL “cealign” function.

**Supp. Figure 10.**
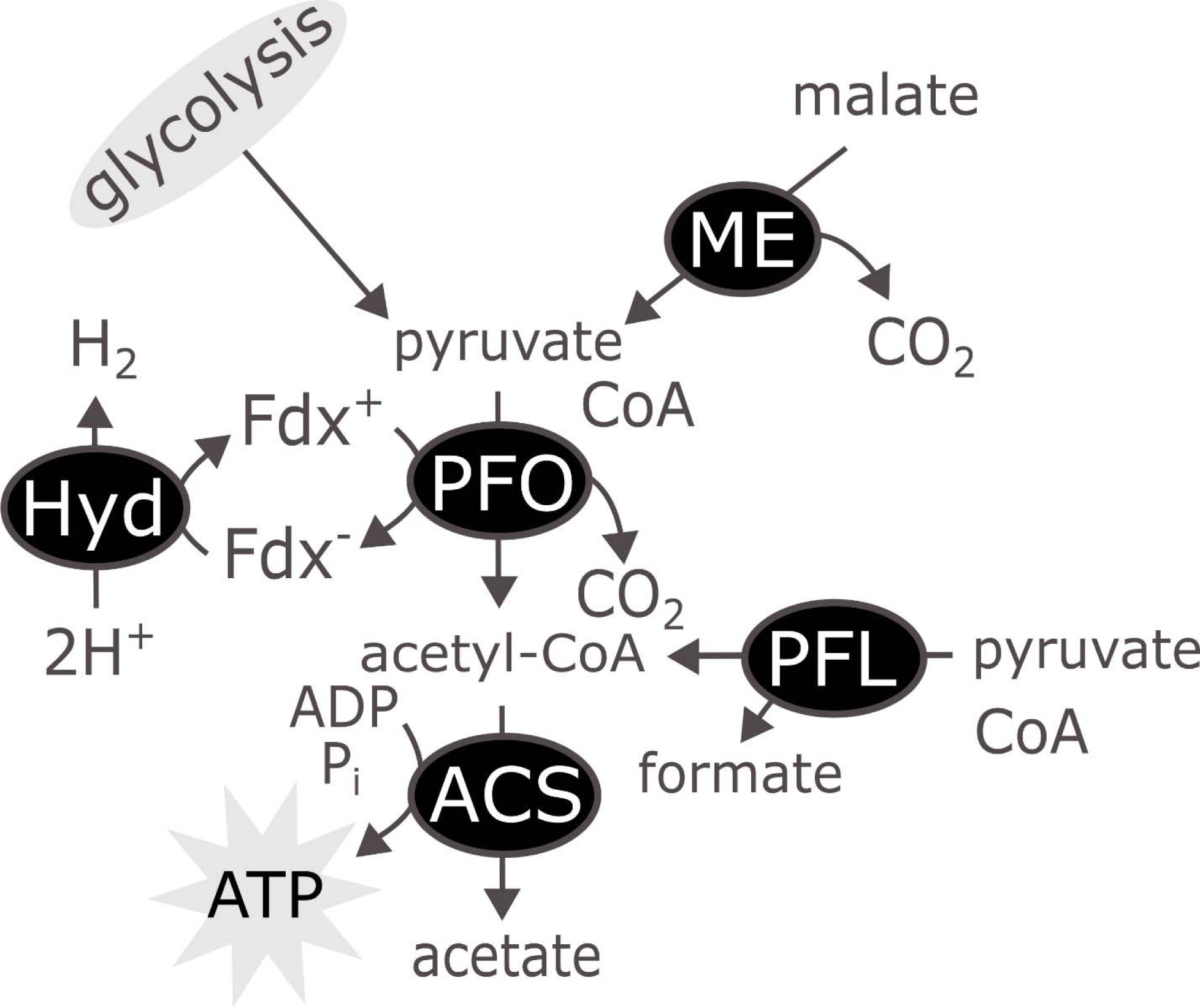
A schematic of the hypothesized ATP production pathway in the BaSk clade. Hyd – Iron-only hydrogenase; ME – Malic enzyme; PFO – Pyruvate:ferredoxin oxidoreductase; ACS – Acetyl-CoA synthase; Fdx – Ferredoxin; PFL – pyruvate-formate lyase.

**Supp. Figure 11.**
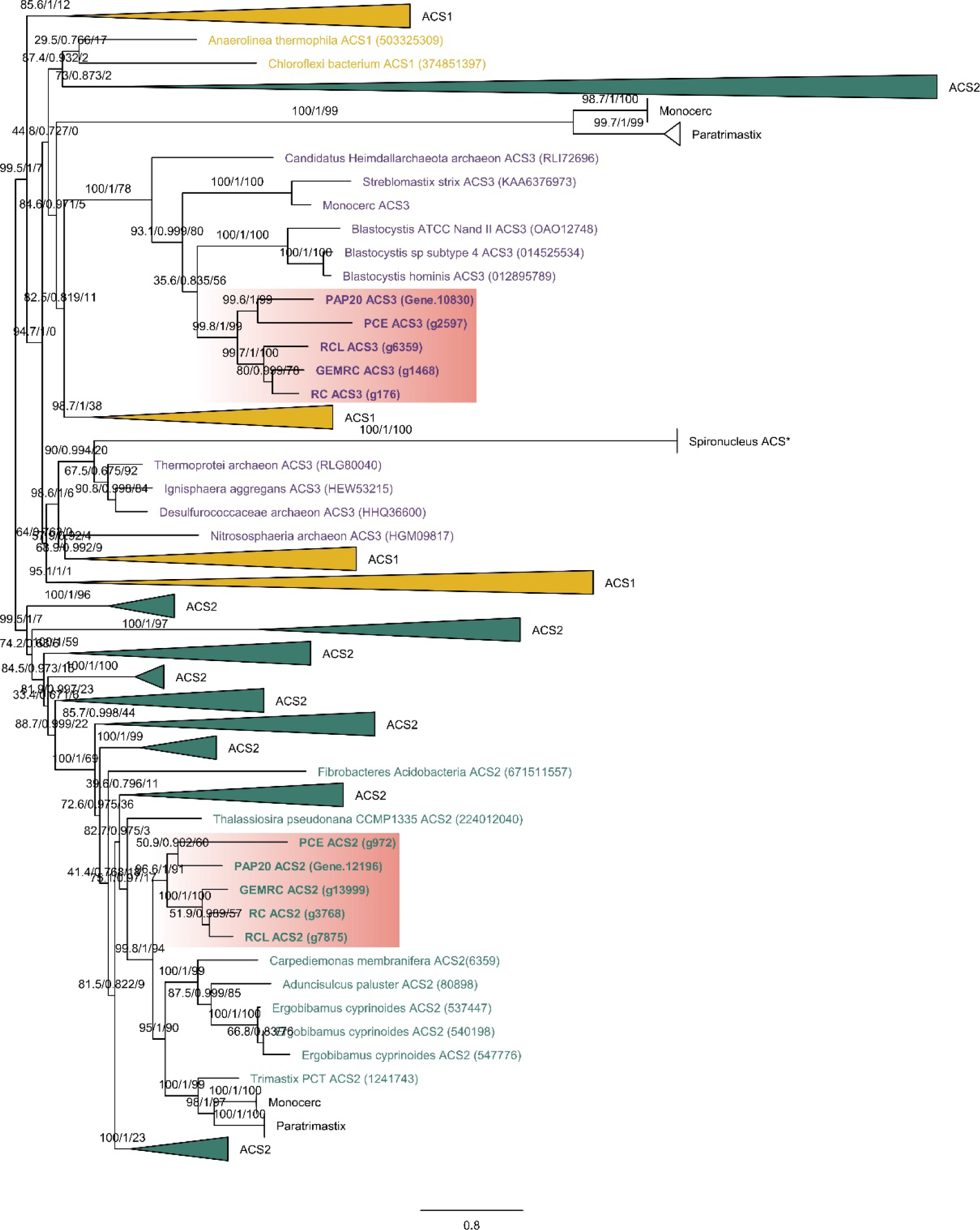
A phylogenetic estimation of the relationship amongst ACS protein sequences in a variety of prokaryotes and eukaryotes, including metamonads. ACS1 is represented in orange, ACS2 in green, and the newly detected ‘ACS3’ in purple. BaSk sequences are bolded and highlighted in red. Major groupings of each ortholog are collapsed into wedges. This phylogeny was constructed using IQ-TREE and the LG+C20+F+Γ model of evolution. Ultrafast bootstrap values are indicated on the branches.

