## Supplementary material for "Extreme mitochondrial reduction in a novel group of free-living metamonads": Figure_legends_2.docx

**Figure 1. ‘BRC’ is a clade of anaerobic protists that branches as sister to all known Fornicata within the Metamonada.** Panels A-D) Differential interference contrast light micrographs of Retortacarp lineages RC (A), GEMRC (B), and RCL (C), as well as *Barthelona* sp. PCE (D) showing overall morphology. Scale bars indicate a length of 10 µm.

Supp. Figure 6. **Estimated phylogenetic relationships amongst a variety of mitochondrial processing peptidases (MPPs) and related metallopeptidases.** This was estimated using IQ-TREE under the LG+C20+F+Γ evolutionary model. Ultrafast bootstrap values are displayed on the branches. BRC sequences are bolded and highlighted in red. Major groups of proteins have been collapsed into wedges.

Supp. Figure 7. **Estimated phylogenetic relationships between members of the mitochondrial carrier family (MCP) with candidate MCP sequences from ‘BRCs’.** Where possible, MCP sequences are labeled with their family subtype designation according to NCBI. This phylogenetic tree was constructed using IQ-TREE and the LG+C20+F+Γ model of evolution. The RC candidate MCP is highlighted in red. Values on the branches represent ultrafast bootstrap values.

Supp. Figure 8 **The phylogeny of Sam50/BamA family in eukaryotes and prokaryotes and predicted structures of the Retortacarp homolog.** A) A phylogenetic reconstruction of the Sam50 candidate proteins in Retortacarp and GEM-Retortacarp, and their relationship to Sam50 (purple) and BamA (green) homolog proteins in eukaryotes and prokaryotes. 8B) Alphafold2 structure prediction using the sequence of the candidate Sam50 protein found in RC 8C) Alphafold2 structure prediction using the sequence of the candidate Sam50 protein found in GEMRC.

Supp. Figure 9 **A schematic of the hypothesized ATP production pathway in the fornifriends.** Hyd – Iron-only hydrogenase; ME – Malic enzyme; PFO – Pyruvate:ferredoxin oxidoreductase; ACS – Acetyl-CoA synthase; Fdx – Ferredoxin

Supp. Figure 10. **A phylogenetic estimation of the relationship amongst ACS protein sequences in a variety of prokaryotes and eukaryotes, including metamonads.** ACS1 is represented in orange, ACS2 in green, and the newly detected ‘ACS3’ in purple. BRC sequences are bolded and highlighted in red. Major groupings of each ortholog are collapsed into wedges. This phylogeny was constructed using IQ-TREE and the LG+C20+F+Γ model of evolution. Ultrafast bootstrap values are indicated on the branches.

Table legends

Table 1. BUSCO scores for members of the BRC clade.

Table 2. Summary statistics for the genome, transcriptome, and predicted proteomes of the BRC clade.

Supp. Table 1. Sequence and additional information for found and queried orthologs mentioned throughout this study.

Supp. Table 2. Found PFO and HydA proteins within members of the BRC clade, with annotation as to additional domains fused to these proteins.
