## Supplementary figures and images for "Extreme mitochondrial reduction in a novel group of free-living metamonads"

### Supp_figure_1_SmsC.pdf

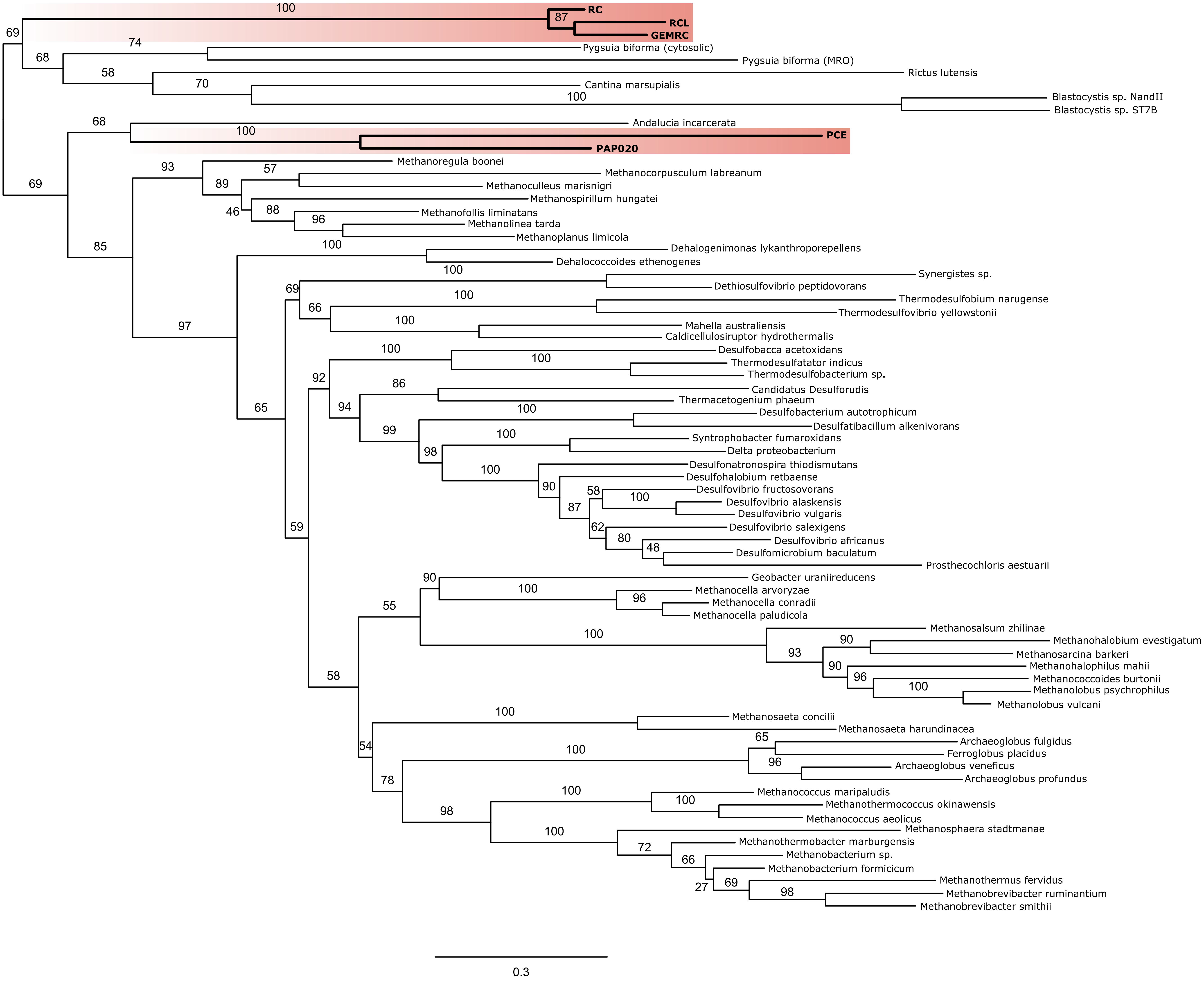

### Supp_figure_2_SmsBD.pdf

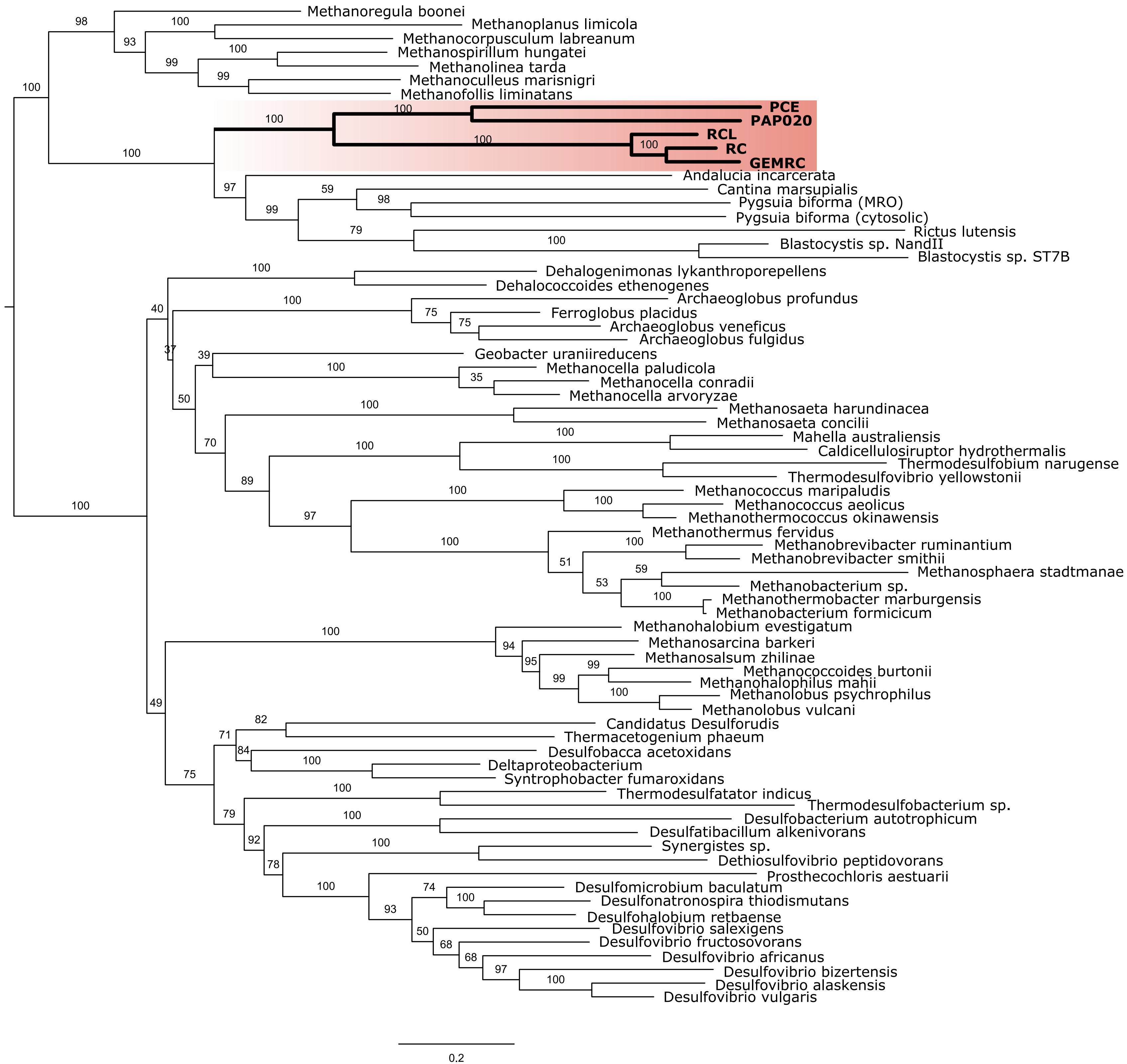

### Supp_figure_3_cysteine_desulfurase.pdf

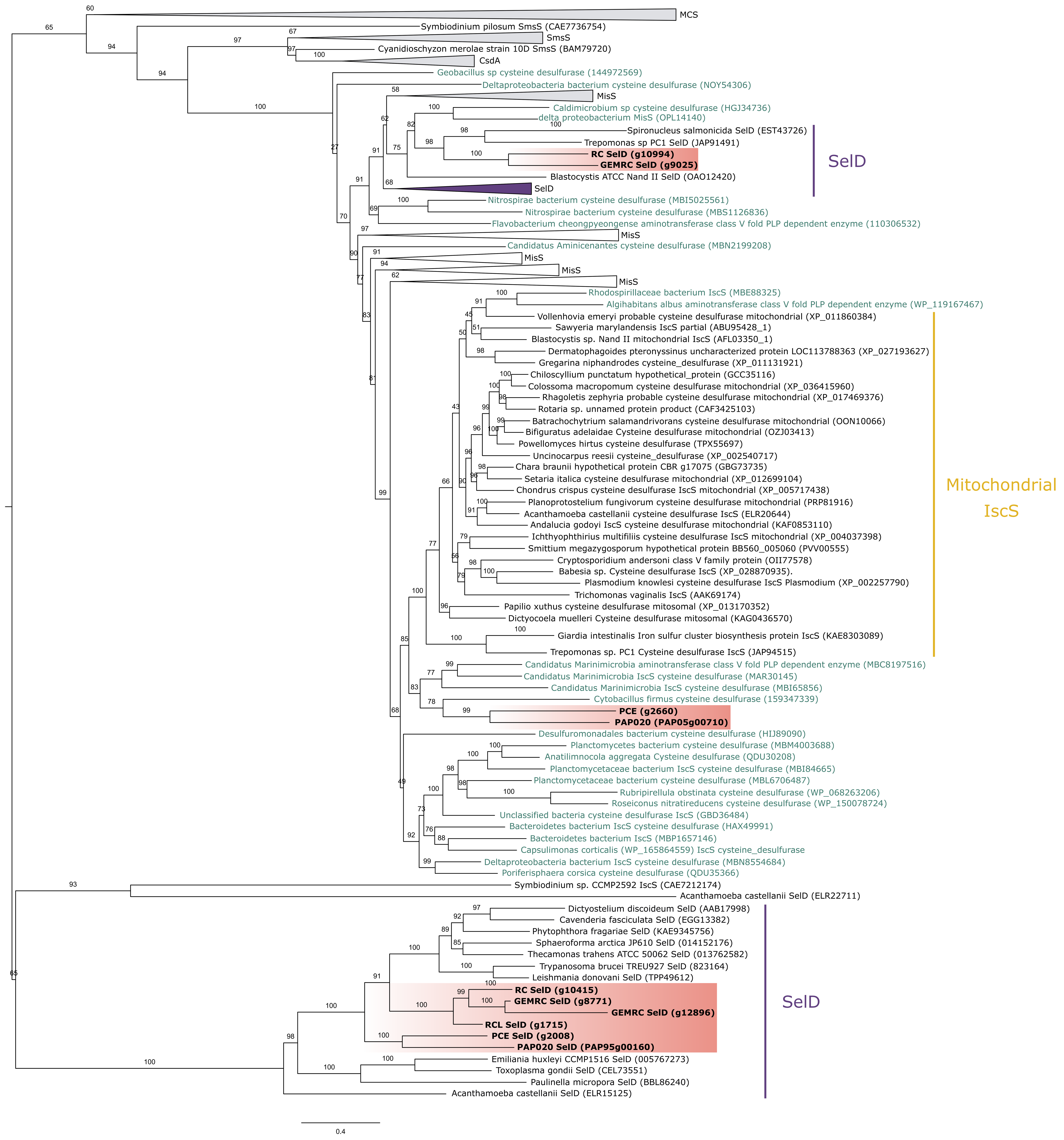

### Supp_figure_4_cpn60.pdf

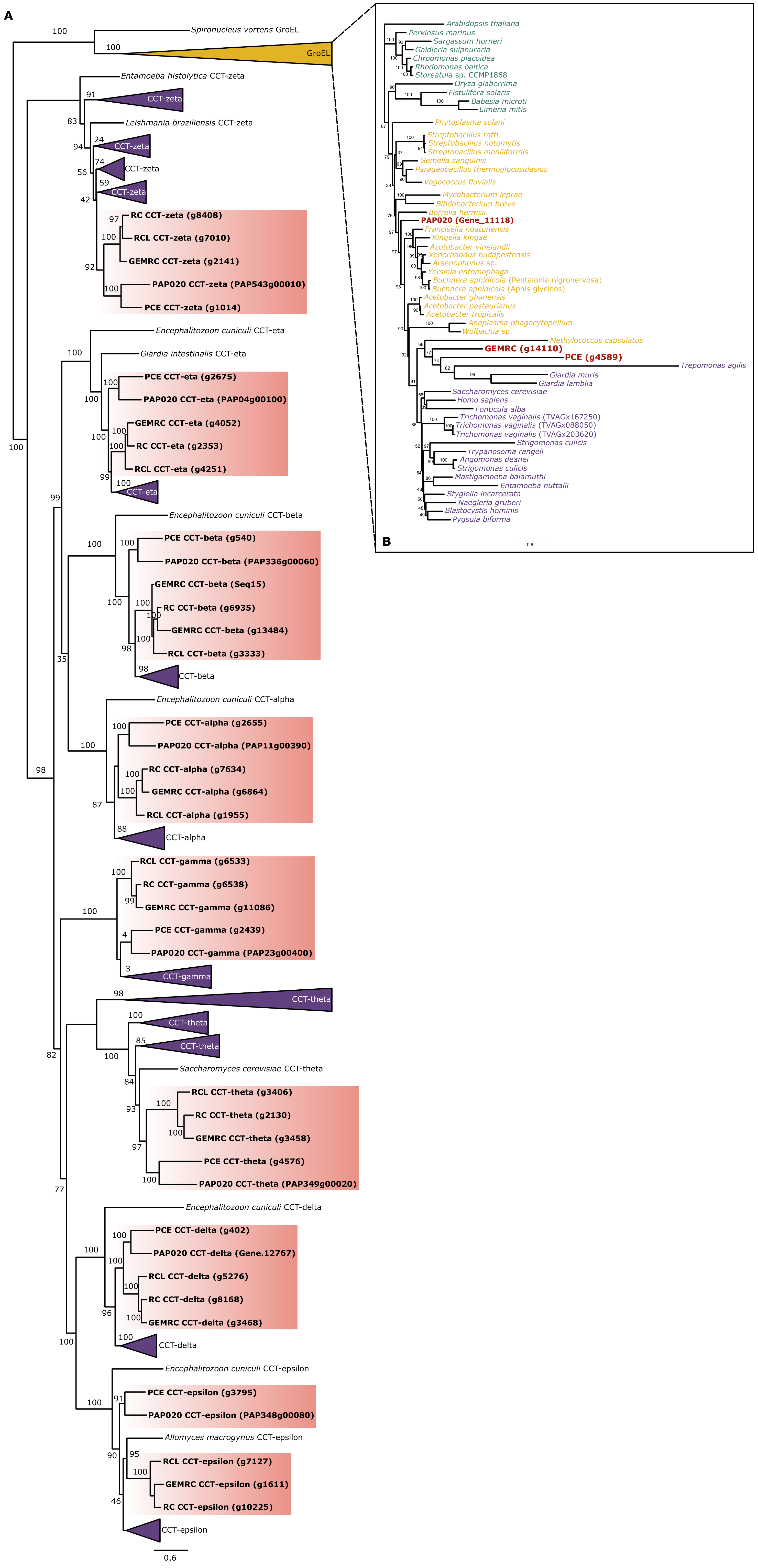

### Supp_figure_5_hsp70.pdf

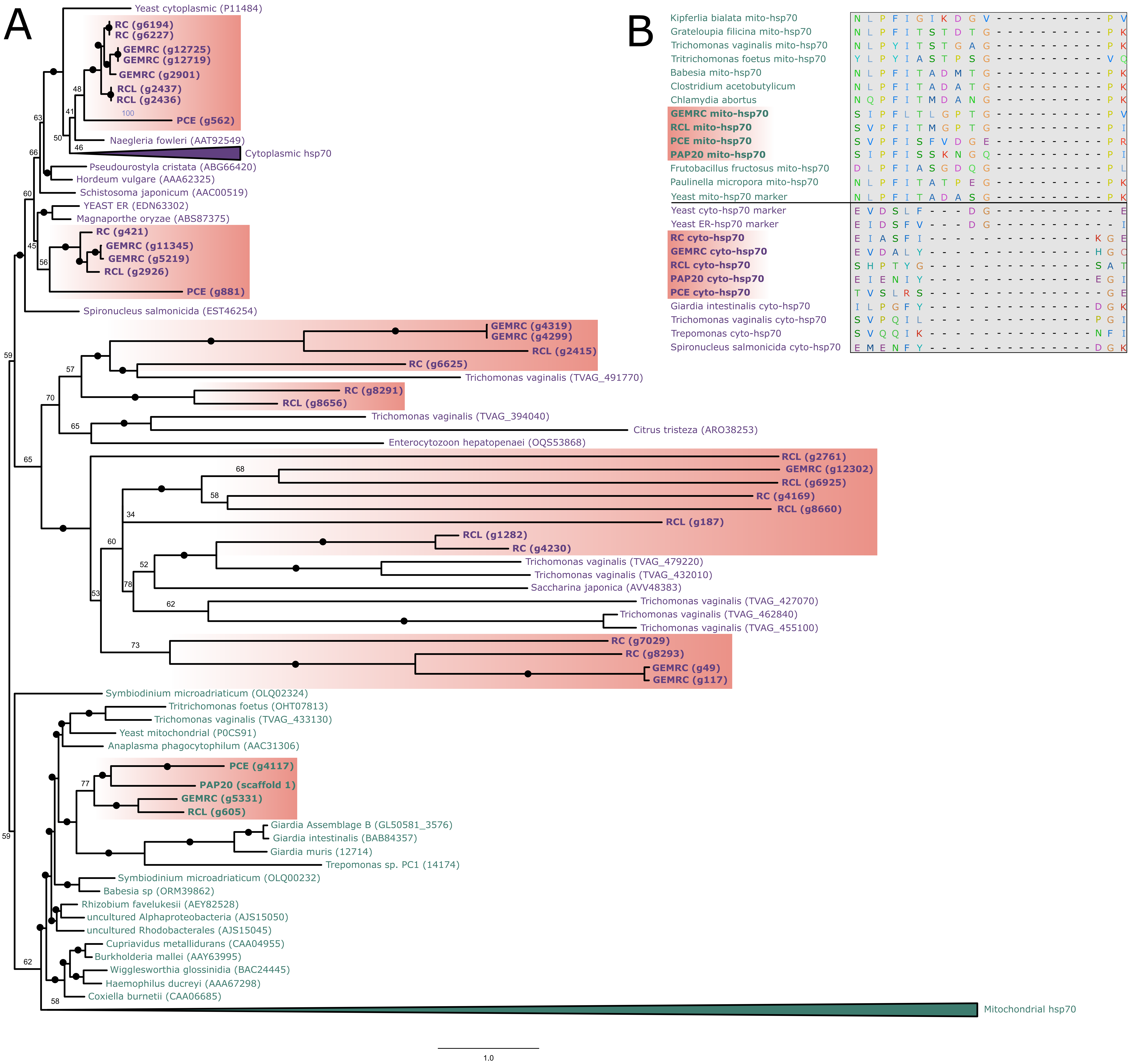

### Supp_figure_6_MPPs.pdf

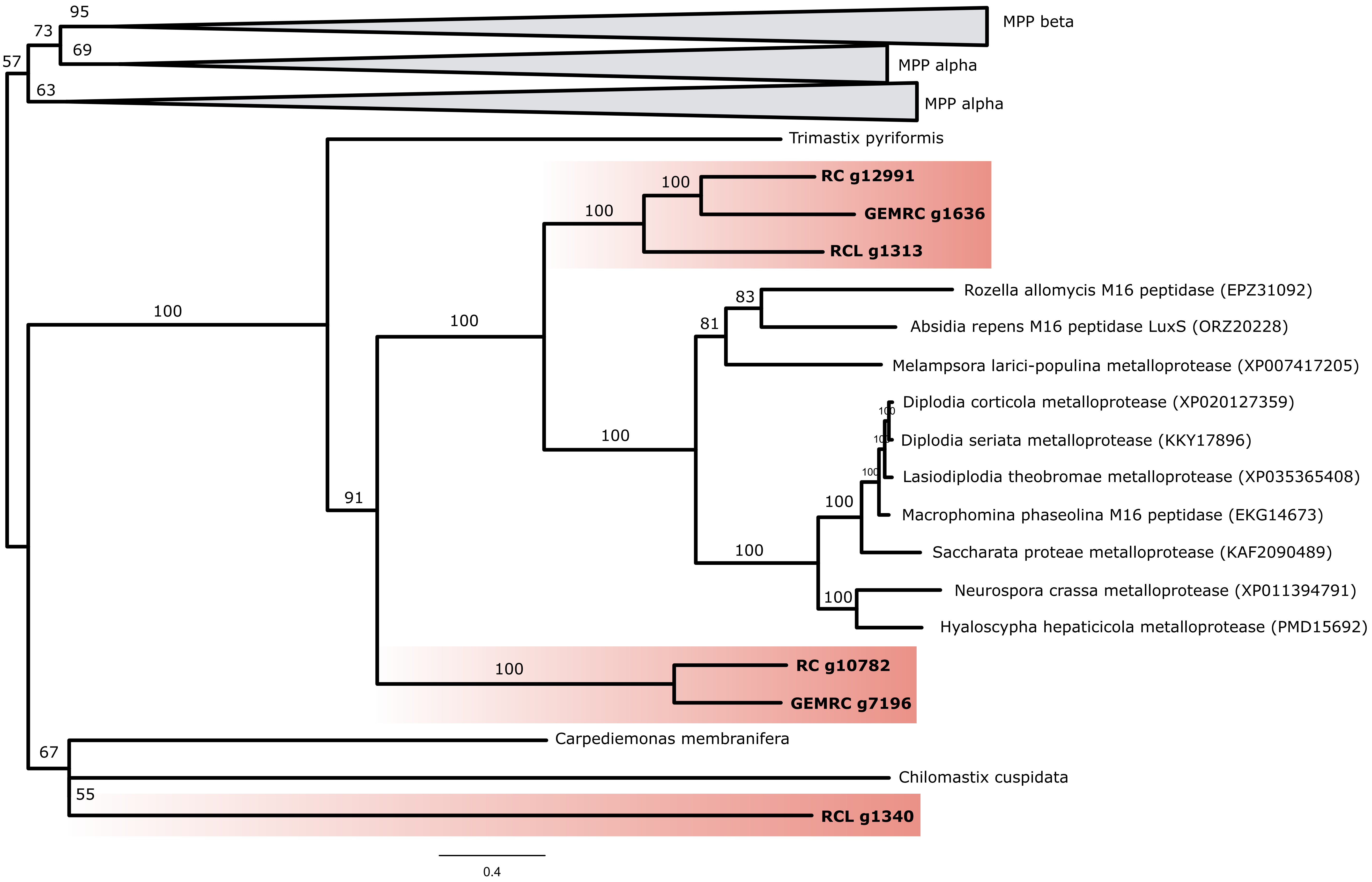

### Supp_figure_7_MCP.pdf

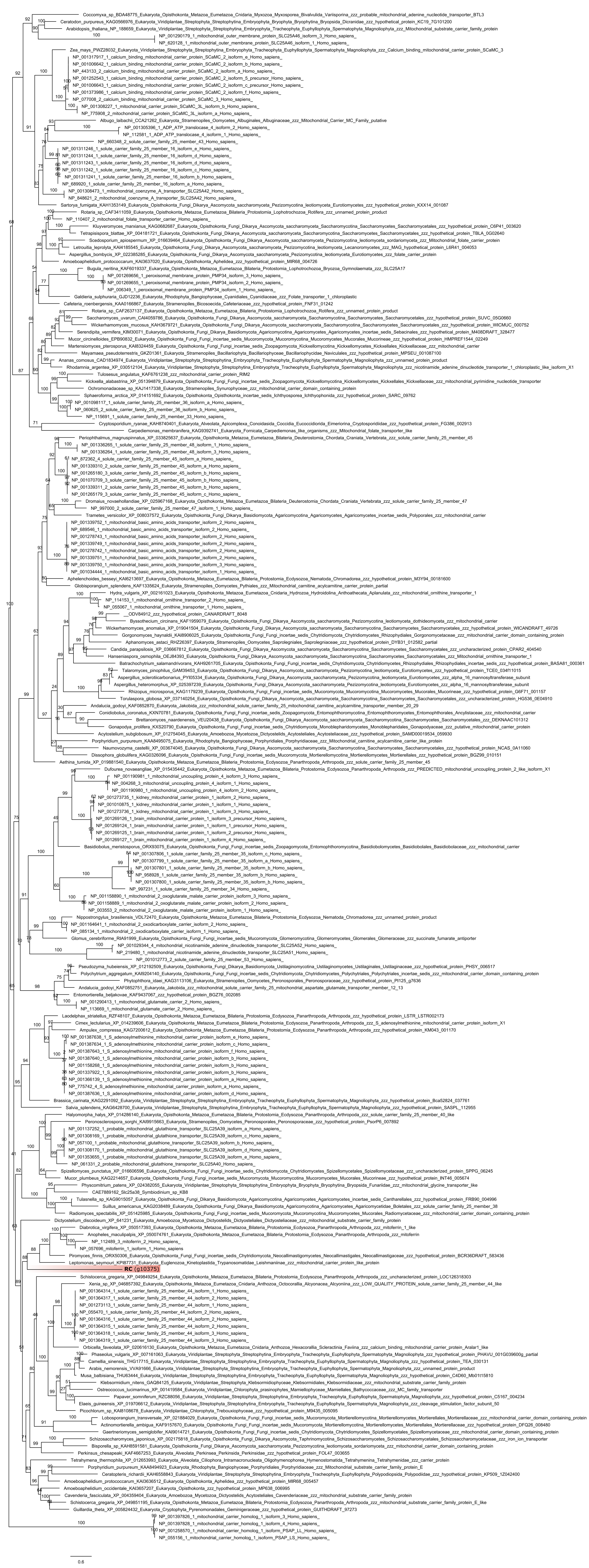

### Supp_figure_8_Sam50.pdf

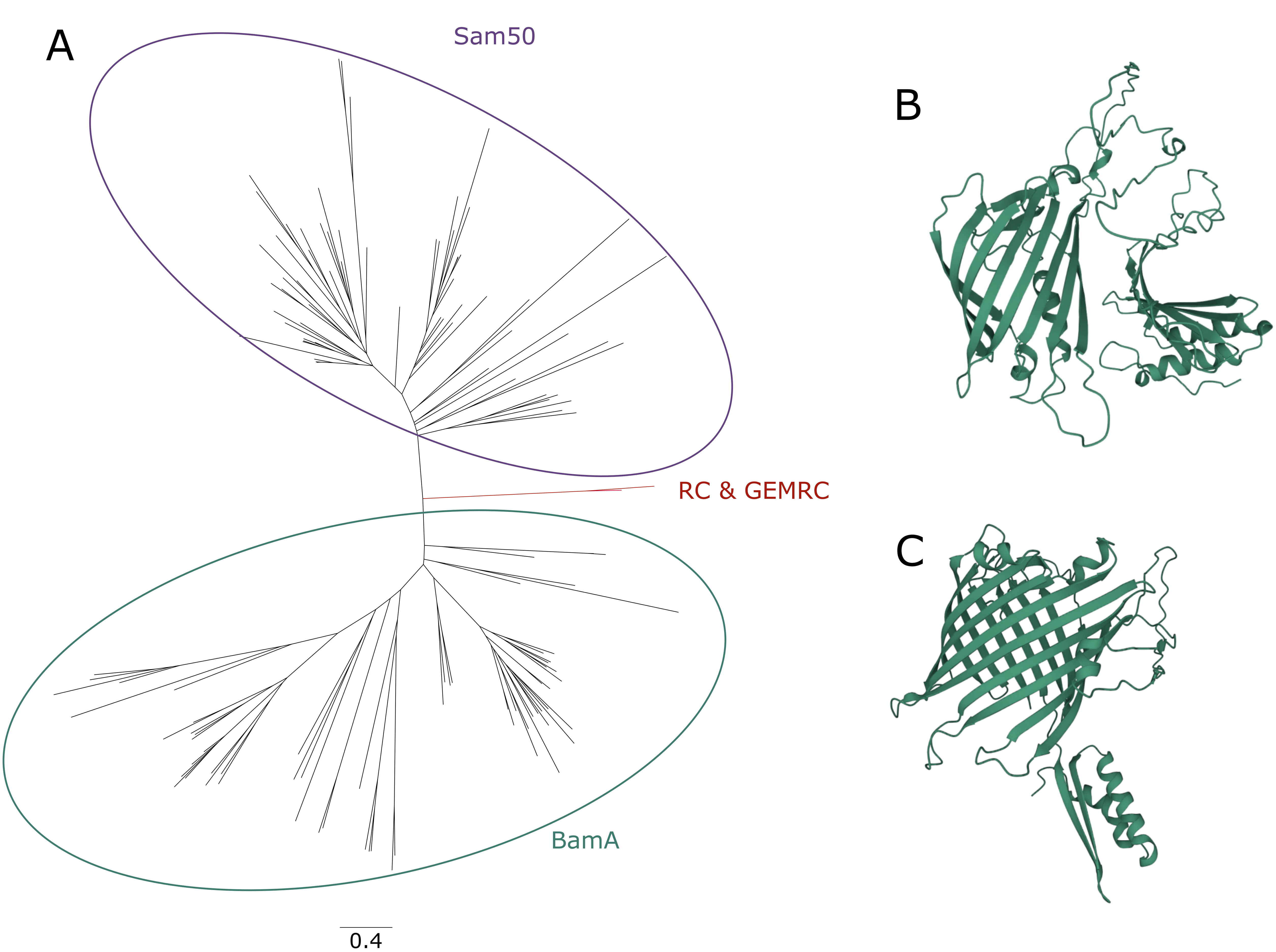

### Supp_figure_9_cyto_metabolism.pdf

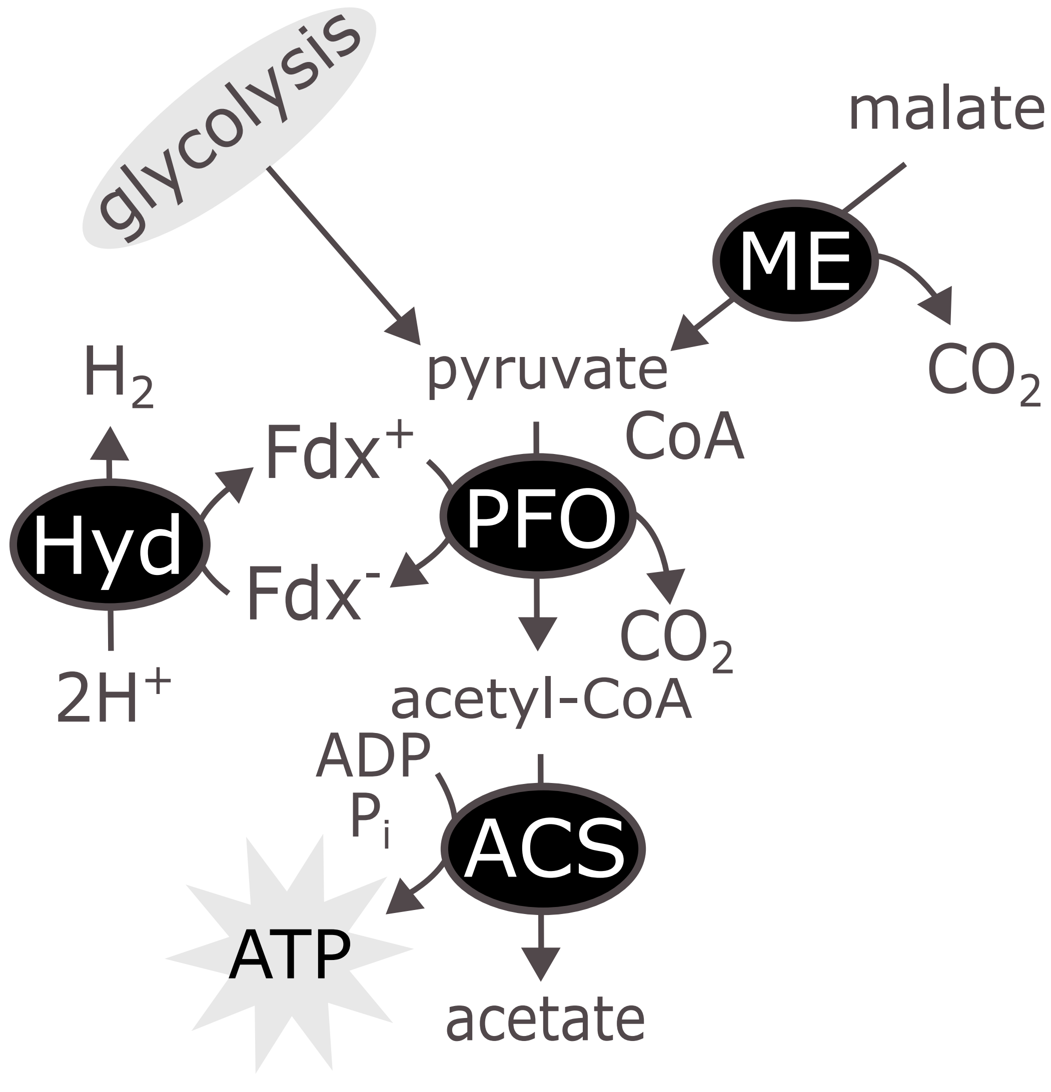

### Supp_figure_10_ACStree.pdf

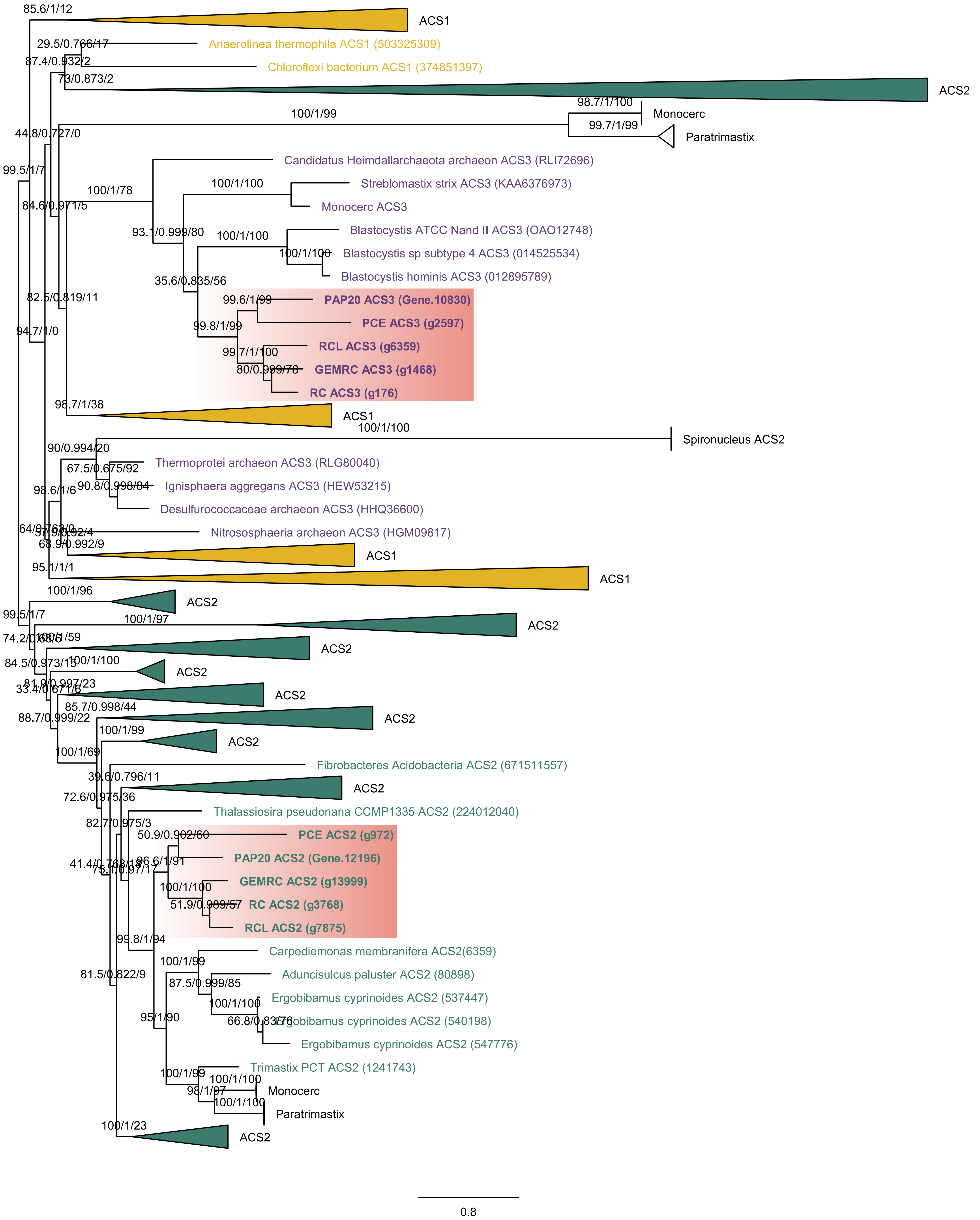
